# Q-MOL: High Fidelity Platform for *In Silico* Drug Discovery and Design

**DOI:** 10.1101/2025.08.06.668254

**Authors:** Anton Cheltsov

## Abstract

For several decades, *in silico* drug discovery methods have held the promise of cheap and efficient ways of streamlining *de novo* discovery of small molecule ligands against novel drug targets. The existing computational methodologies proved to be successful in those cases when a protein target is a relatively rigid molecule (e.g., many enzymes), or when an *in silico* methodology is applied during hit-to-lead optimization of an already potent binder. However, these methods fail when applied to highly flexible or intrinsically disordered protein targets which represent the bulk of pharmacologically important proteins. The major factors contributing to the failure are either incorrect or simplified treatment of protein flexibility, inadequate or overfitted scoring function, absence of correctly annotated druggable pockets, and lack of structural data about potential allosteric sites. The Q-MOL drug discovery platform addresses the problem of treatment of protein flexibility during the course of protein-ligand docking by computationally implementing some of postulates of the energy landscape theory of protein folding. The Q-MOL protein-ligand docking method was thoroughly validated in academic and commercial settings across a panel of more than 60 diverse proteins (of viral, bacterial and animal origin). This methodology, originally developed for high fidelity virtual ligand screening, is also used for the prediction of ligand binding sites. The ability to reliably predict ligand binding sites on the surface of novel and unannotated protein targets is critical for successful drug discovery. The Q-MOL drug discovery platform is agnostic to the nature of a protein target. Rigid, flexible or intrinsically disordered proteins are processed by the same protein-ligand docking protocol. Importantly, the software does not require the presence of structurally well-defined binding pockets or sites. The correct and validated implicit treatment of protein flexibility allows for successful discovery of small molecule ligands against allosteric sites lacking any structural features of a classical druggable pocket. To demonstrate the power of its methodology, Q-MOL was used to detect ligand binding sites on the surface of the bHLHZ domain of the c-Myc protein. Q-MOL virtual ligand screening was then used to dock a human metabolites library against one of the identified binding sites. The top hits were cross-referenced against published data, notably identifying dehydroascorbic acid as one of the top-ranked compounds. In contrast to other docking programs, all of the Q-MOL *in silico* drug discovery features were thoroughly validated *in vitro*, *in cellulo* and *in vivo* when possible. In this paper, the results of several previously published drug discovery projects are briefly presented and related to relevant computational features of the software. The corresponding protein targets include viral proteinases (West Nile, Hepatitis C, Dengue and Zika viruses), hemopexin domain of membrane type I matrix metalloproteinase, retinoid X receptor α, and β-catenin armadillo repeat domain. It has been also shown that contrary to existing belief, highly flexible proteins (or their domains) represent significantly more amenable drug discovery targets than rigid proteins. Flexible and intrinsically disordered proteins exist in a vast array of environment-dependent conformational states, and thus, they can accommodate a significantly larger number of ligand chemotypes ensuring both potency and specificity of interactions with a protein target. It has been also demonstrated that Q-MOL binding sites prediction and ligand docking methodologies, initially developed for protein targets, can be successfully applied "As Is" to polynucleotide-based structures, such as non-coding viral RNA molecules.

## Introduction

Drug discovery and development are challenging high risk-high reward endeavors. Initial identification of new drug leads for a new drug target is both an essential and critical step in any drug discovery/drug development process. Because of the low cost and high speed of the computations, *in silico* drug discovery methodology such as protein-ligand docking, a core of virtual ligand screening (VLS) technology, has always been of great interest to the pharmaceutical industry. However, despite decades of financial and labor investments into research and development only relatively modest results have been achieved. Specifically, the bulk of most pharmaceutically lucrative protein targets still remains virtually unreachable by *in silico* methods. These protein targets are also hard to approach with more conventional methods such as high throughput screening (HTS). These proteins belong to the families of transcription factors, adaptor and scaffolding proteins, allosteric modulators of larger complexes, regulatory domains of certain enzymes and many others. One property, these proteins share, is high flexibility or intrinsic disorder of their structure. These are also known as intrinsically disordered proteins (IDPs). The high degree of structural flexibility makes them also very difficult to work with using wet chemistry methods.

As expected, available VLS methods perform well when applied to rigid enzyme targets or to the virtual benchmarks that are composed of assorted families of protein targets. In both cases, the protein structure remains largely fixed in terms of backbone conformational changes, and ligand binding sites exist as well-defined druggable pockets. Thus, the success of existing computational methods with rigid protein targets is largely a function of mere surface and chemical complementarity of a ligand and a binding pocket. The direct application of these methods to novel flexible protein targets results in an intractably large number of false positives. It is well understood that the correct treatment of protein flexibility must be incorporated into protein-ligand docking methods in order to apply them successfully to flexible and intrinsically disordered proteins.

The treatment of protein flexibility is an essential problem to solve for any protein-ligand docking method to yield accurate results. However, there is another equally important challenge that needs to be addressed. It is the problem of ligand binding site prediction. The correct prediction of ligand binding sites is critical for protein annotation and drug discovery. A number of ligand binding sites prediction methods exist^1^. These methods employ either statistics or machine learning techniques using geometric or energy features of known liganded hollows and cavities as input. However, since the ligand binding sites are also influenced by protein flexibility, the application of such methods to flexible proteins results in large number of false positives ^1^. In addition, the large number of non-enzyme proteins of known structure do not contain classical ligand binding pockets within their structures. Instead, they appear to use extensive surface interfaces to interact with other proteins. It is also understood that many of such proteins contain allosteric and cryptic binding sites that are used for functional modulation. These ligand binding sites are created on the fly as a result of specific conformational transitions of the protein of interest. Thus, the ligand binding site prediction methods must incorporate the correct treatment of protein flexibility to produce sensible results.

This paper presents novel protein-ligand docking and ligand binding site(s) prediction methodologies that address protein flexibility *via* efficient computational implementation of general concepts of the energy landscape theory of protein folding ^2^. It has been demonstrated that the methodology can be successfully applied to highly flexible proteins, IDPs, and pocketless ligand binding sites. It has been also demonstrated that all of the Q-MOL features, initially developed for protein targets, can be applied "As Is" to non-protein molecules such as polynucleotides (e.g. non-coding viral RNA molecules). The theoretical concepts and validity of certain implementation details are validated here by *in silico*, *in vitro, in cellulo* and *in vivo* results of the application of the Q-MOL VLS and the ligand binding site(s) prediction method.

## Results and Discussion

The Q-MOL platform was validated in over 60 *in vitro, in cellulo* and *in vivo* (when possible) drug discovery projects dealing with either novel protein targets, or novel inhibition mechanisms of known protein targets. All biological assays were performed by independent scientists within collaborating laboratories. The results of several thoroughly validated drug discovery applications of Q-MOL platform are presented below.

*West Nile Virus (WNV) NS2B-NS3pro Flavivirus 2-component Processing Proteinase.* There were several, relatively successful, high throughput screening (HTS) attempts to identify inhibitors of WNV NS2B-NS3pro ^3–8^. Previous HTS studies suggested that the 5-amino-1-(phenyl)sulfonyl-pyrazol-3-yl class inhibitors interacted with the NS2B-binding cavity in the NS3pro domain and that they interfered with the unique feature of the flaviviral proteinases such as the productive interactions of the NS2B cofactor with the NS3pro domain^3, 4^. In turn, the enzyme active site is largely conserved in the human and viral serine proteinases, and it lacks the structural features, which could be readily exploited to achieving both the specificity and potency of the inhibitors. Thus, small molecule ligand interference with the productive conformation of the NS2B cofactor represents a superior drug discovery strategy when compared to targeting of the active site of the viral proteinase (Fig. 1). To validate this hypothesis, the Q-MOL VLS technology was used to identify *de novo* allosteric small molecule inhibitors of NS2B-NS3pro (Fig. 2)^9^. Note that only 50 ligands were assayed *in vitro* to identify 18 active compounds (36% success rate), while Q-MOL primary VLS converged to only 85 ligands out of 275,000 (99.97% enrichment). Out of 18 active compounds, 3 inhibitors were in nanomolar range, and 2 in low micromolar range.

**Figure 1.**
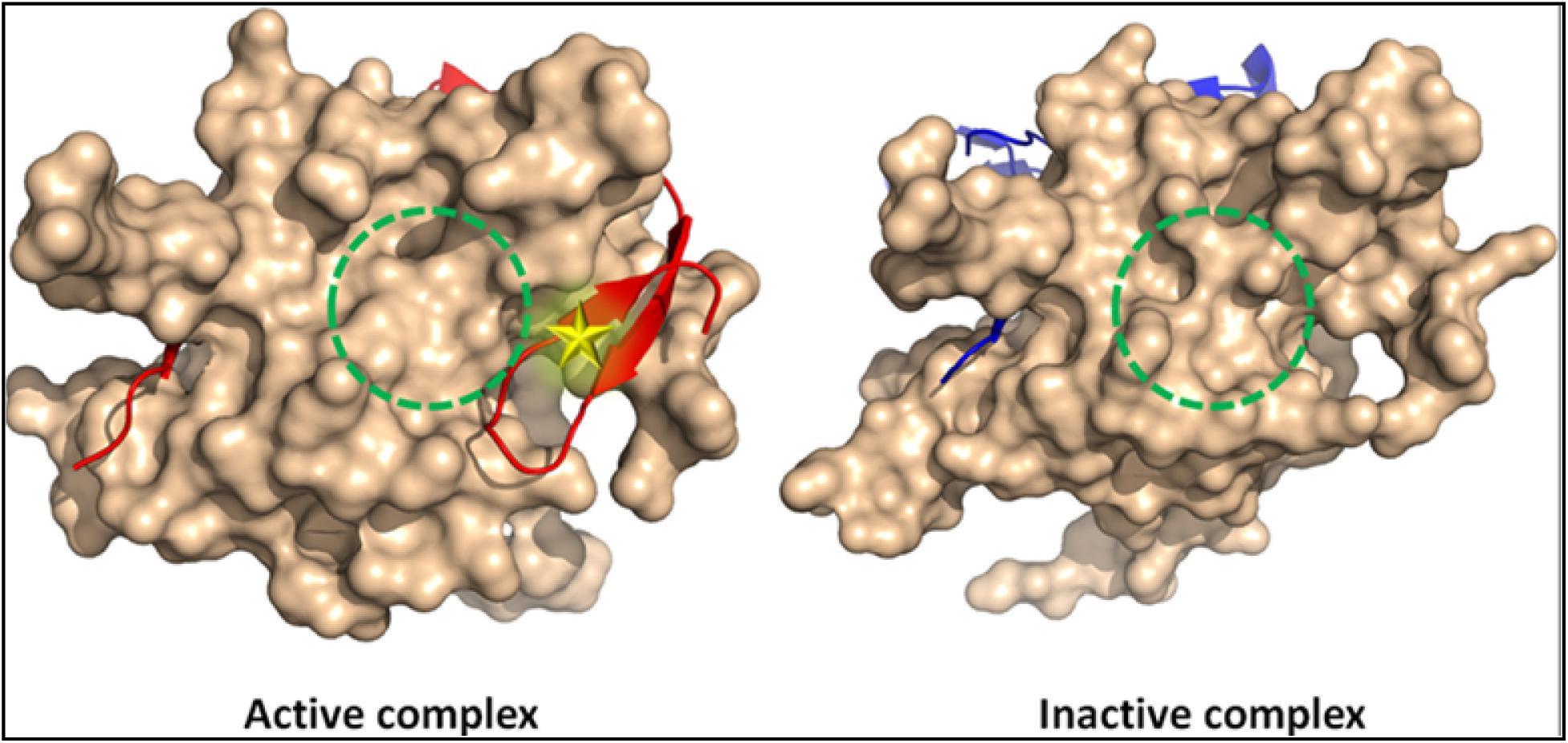
Strategy of discovery of allosteric small molecule inhibitors of WNV NS2B-NS3pro via interference with NS2B peptide cofactor placement. WNV NS2B-NS3pro is shown as molecular surface, location of the docking site is depicted as yellow star, location of the active site is shown with green circle. NS2B cofactor is displayed in red in active complex, and in blue in inactive complex.

**Figure 2.**
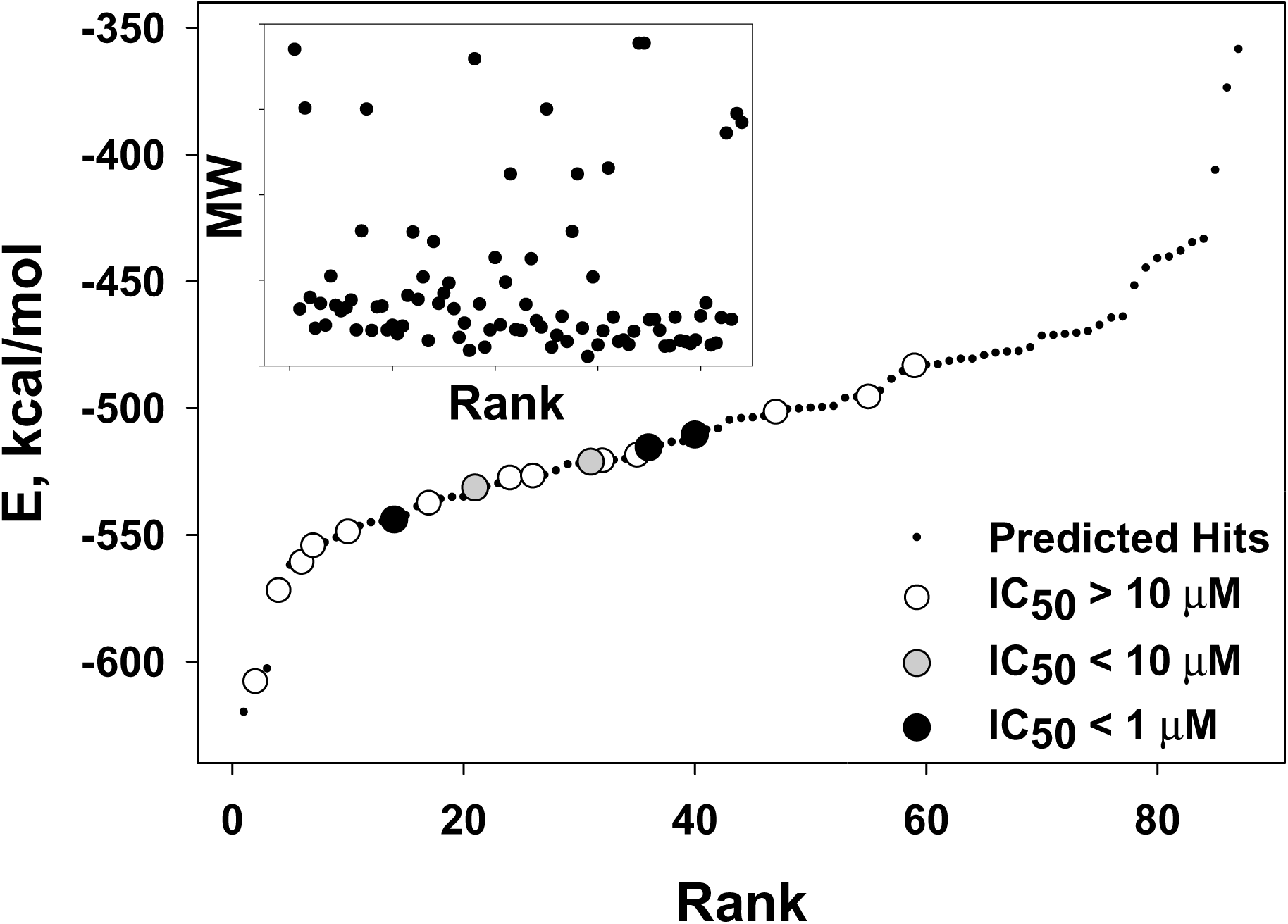
Primary virtual ligand screening (VLS) and *in vitro* validation of allosteric small molecule inhibitors of WNV NS2B-NS3pro proteinase. The complete NCI DTP SDF library was used as a source of ligands (≈ 275,000 structures). The Q-MOL primary VLS converged to 85 ligands (99.97% enrichment factor), out of which, top 50 predicted binders were ordered from NCI and tested in *in vitro* cleavage assay. Out of 50 assayed ligands, 3 ligands had IC_50_ < 1 µM (black filled circles), 2 ligands had IC_50_ in range 1 µM - 10 µM, and 13 ligands had IC_50_ > 10 µM (18 active hits in total, 36% success rate). Inset: plot of ligands molecular weight (MW) vs. their rank.

### Hepatitis C virus (HCV) NS3 serine proteinase (NS3/4A)

NS3/4A is a chemotrypsin-like NS3 serine proteinase that cleaves the viral polyprotein precursor at the NS3/NS4A, NS4A/NS4B, NS4B/NS5A and NS5A/NS5B junction regions. The individual NS3/4A, however, is inactive. For its cleavage activity *in vitro* and *in vivo*, NS3/4A requires either the full-length NS4A co-factor or, at least, its 14-residue hydrophilic central portion ^10, 11^. Because the NS3/4A mechanism of activation is identical to that of WNV NS2B-NS3 two-component proteinase, the same *in silico* drug discovery strategy of targeting several allosteric sites of interaction of NS3/4A proteinase with its NS4A cofactor was successfully applied (Fig. 3) ^12^. Note that the Q-MOL protein-ligand docking was successful in finding biologically active ligands for all 3 targeted allosteric sites. The primary Q-MOL VLS converged to 84, 87 and 88 top individual hits for sites 1, 2 and 3, respectively. After visual inspection, 14, 37 and 36 ligands were selected for ordering from NCI DTP for sites 1, 2 and 3, respectively. Out of which, 7, 15 and 18 ligands exhibited activity for sites 1, 2 and 3, respectively (Fig. 3). Docking Site 3, structurally similar to WNV NS2B-NS3 docking sites, produced the largest number of biologically active potent small molecule ligands. The validated hits obtained for Site 3 after primary Q-MOL VLS were further computationally optimized, and new analogues re-tested (Fig. 4) (see Computational Optimization of Primary VLS Hits in Methods). As a result, the most potent of identified inhibitors for Site 3, NSC704342 had wild-type HCV NS3/4A IC_50_ = 183 nM (compare to telaprevir IC_50_ = 148 nM). NSC704342 also maintained its potency across the panel of most common HCV NS3/4A mutations ^12^.

**Figure 3.**
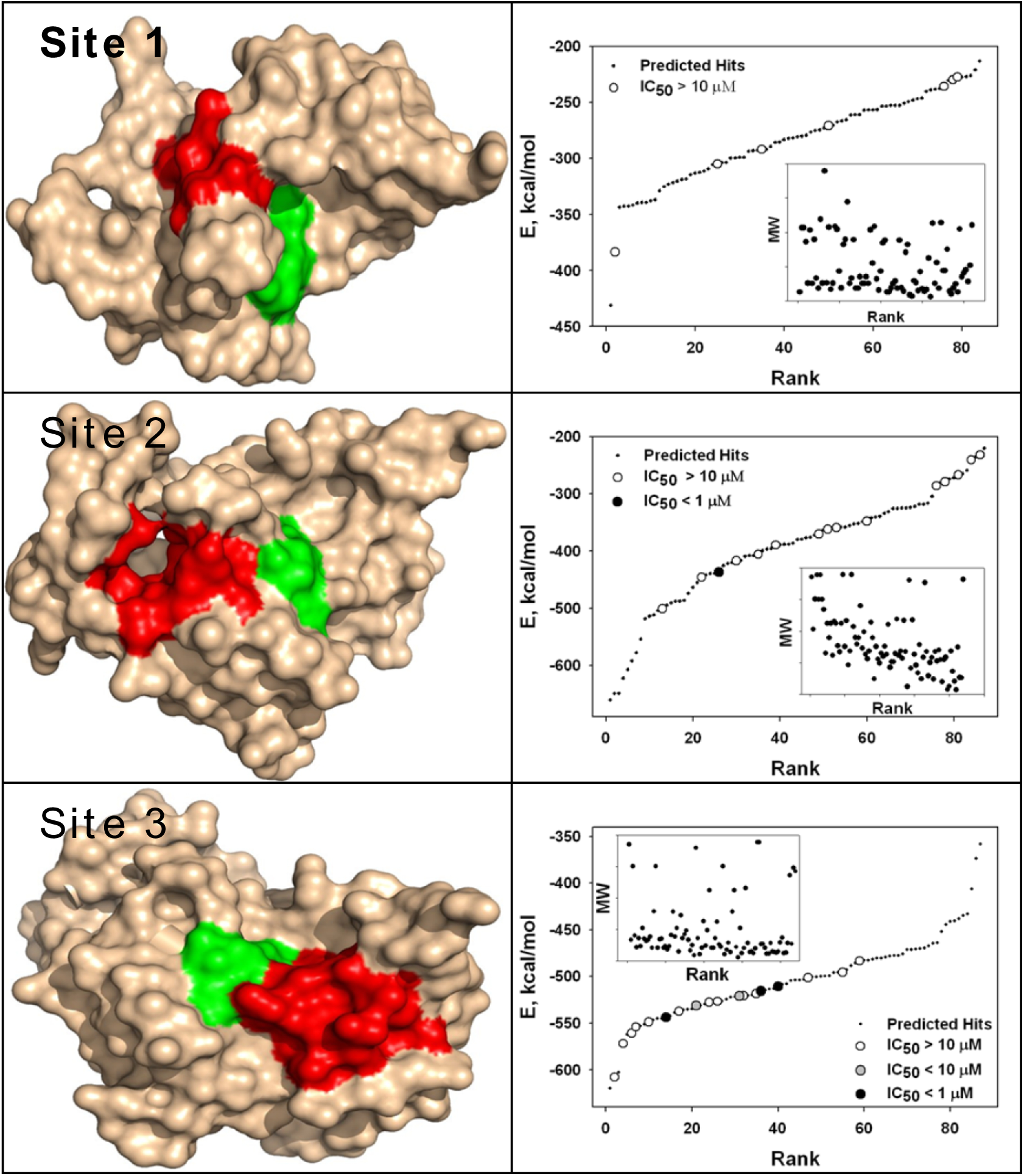
HCV NS3/NS4A proteinase docking sites selection, primary VLS results and *in vitro* validation of allosteric small molecule inhibitors. The complete NCI DTP SDF library was used as a source of ligands (≈ 275,000 structures). The primary Q-MOL VLS converged to 84, 87 and 88 top individual hits for sites 1, 2 and 3, respectively. After visual inspection, 14, 37 and 36 ligands were selected for ordering from NCI DTP for sites 1, 2 and 3, respectively. Out of which, 7, 15 and 18 ligands exhibited activity for sites 1, 2 and 3, respectively. The location of docking sites are displayed with red molecular surface patch, green molecular surface patch corresponds to the active site of the enzyme. Inset: plot of ligands molecular weight (MW) vs. their rank.

**Figure 4.**
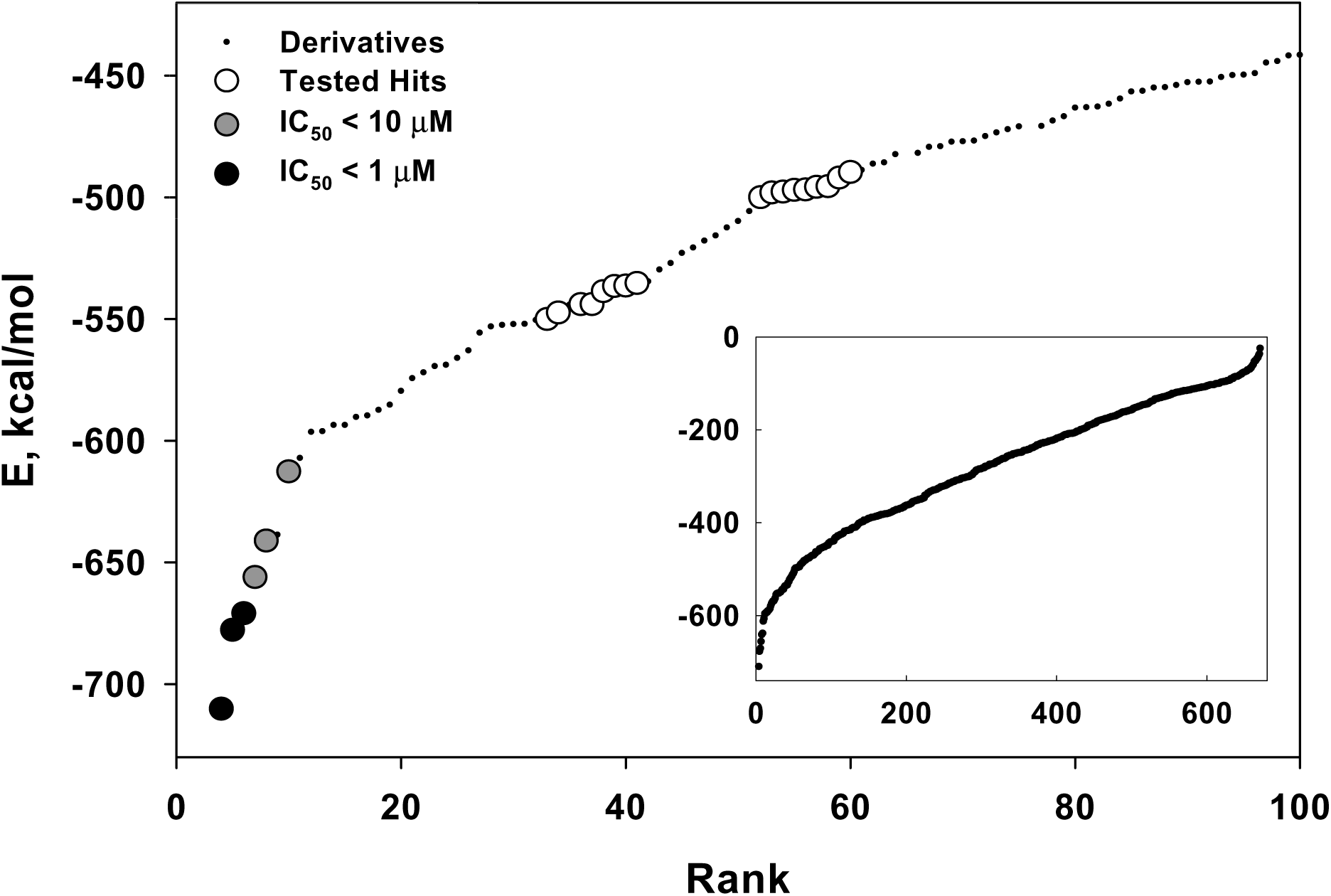
*In silico* structure-activity-relationship optimization of primary VLS hits. Total of 665 analogs of primary VLS hits were selected from database based on Q-MOL chemical fingerprint distance. The new ligands were docked into the Site 3 (Fig. 3) of the HCV NS3 proteinase (black dots, inset). Out of which, 23 ligands were obtained and tested *in vitro* (white-filled circles). Out of 23 tested ligands, 3 ligands had *IC_50_* values below 1 μM (black-filled circles), and another 3 ligands below 10 μM (grey-filled circles). *E*, relative binding energy; *Rank*, ligands ranking after sorting by relative binding energy from lowest (best) to highest (worst).

### Zika virus (ZIKV) and Dengue virus (DENV) serine proteinases

ZIKV and DENV virus belong to the same family of flaviviruses as the WNV virus. The N-terminal region of NS3 protein of both ZIKV and DENV viruses encode a serine proteinase, while the C-terminal region encodes proteinase peptide cofactor NS2B. ZIKV and DENV NS2B-NS3 proteinase are structurally very similar to WNV proteinase complex. Thus, it was hypothesized that the same allosteric mechanism of inhibition as for WNV proteinase can be exploited, and previously identified WNV NS2B-NS3 allosteric inhibitors should be able to inhibit both ZIKV and DENV NS2-NS3 proteinases. Indeed, this hypothesis was successfully validated, and several allosteric WNV proteinase inhibitors were also able to inhibit ZIKV and DENV proteinase at low micromolar concentrations ^13, 14^. The most potent WNV and ZIKV NS2B-NS3 allosteric inhibitor NSC86314 (WNV IC_50_ = 260 nM) inhibited ZIKV proteinase at IC_50_ = 1.12 µM. Finally, the pan flaviviral proteinase inhibitor NSC86314 has been shown to be active *in vivo* in an independent study^13^, and it was co-crystallized with ZIKV NS2B-NS3-Mut7 proteinase (PDB 7M1V)(Fig. 4) ^14^. All together, these results provided the ultimate proof of the approach and validation of the Q-MOL protein-ligand docking methodology.

### Retinoid X Receptor α (RXRα) and Hemopexin (PEX) Domain of Membrane Type 1 Matrix Metalloproteinase (MT1-MMP)

The Q-MOL protein-ligand docking methodology was further validated on non-enzyme target proteins such as receptor RXRα ^15^, and PEX domain of MT1-MMP ^16^. RXRα plays an important role in tumorigenesis, and some of its ligands possess potent anticancer activities. A novel RXRα antagonist NSC640358 was identified using the Q-MOL primary VLS. The compound acted as selective RXRα ligand and promoted TNFα-mediated apoptosis of cancer cells. The site-directed mutagenesis experiments indicated that NSC640358 binds to RXRα in a mode distinct from other known receptor antagonists.

Targeting of MT1-MMP PEX domain represented a novel approach to safe and specific inhibition of matrix metalloproteinase. PEX domain homodimerization is required for MT1-MMP translocation to the surface of a cancer cell where it facilitates metastasis. Thus, modulating a specific extracellular function of MT1-MMP via interference with PEX domain homodimerization is potentially a safe way to impact metastasis of certain common cancers such as breast cancer without interfering with its critical intracellular functions. The Q-MOL platform was successfully used to discover small molecule modulator of PEX domain that exhibited significant activity *in vitro* and *in vivo* (Fig. 5)^16^. After a single round of the Q-MOL primary VLS, only 19 ligands were screened *in vitro* to discover a potent hit that was also active *in vivo*.

**Figure 5.**
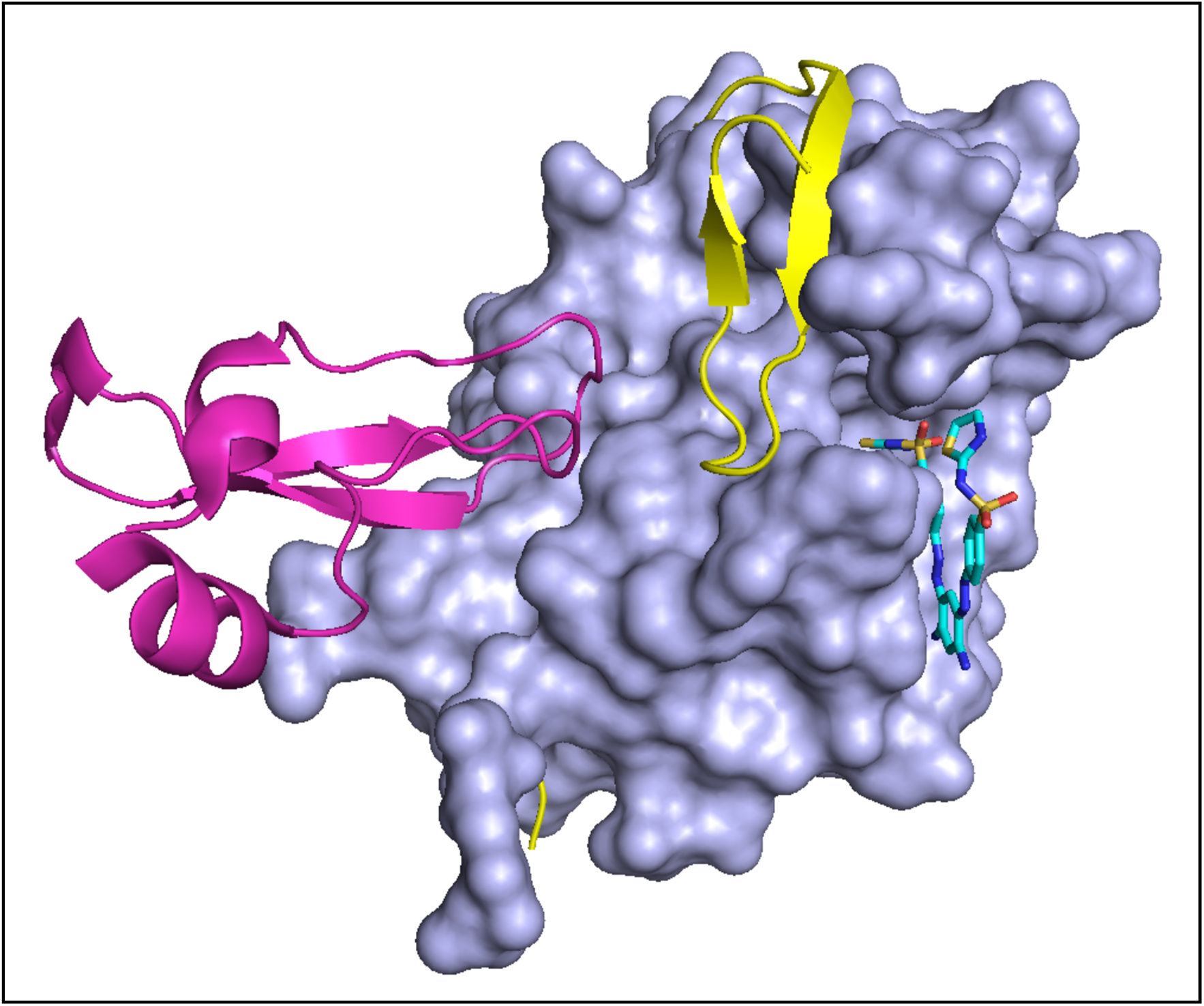
Crystal structure of Zika virus NS2B-NS3 protease mutant in complex with the compound NSC86314 in the super-open conformation (PDB 7M1V). The chain A of Zika virus NS2B-NS3 protease is shown as molecular surface (pale blue), co-crystallized ligand NSC83614 is shown in stick style. Crystal Structure of the West Nile virus NS2B-NS3 protease complexed with bovine pancreatic trypsin inhibitor (PDB 2IJO) was superimposed with the chain A of Zika virus NS2B-NS3 protease. Pancreatic trypsin inhibitor is shown in purple, WNV NS2B cofactor is shown in yellow.

**Figure 6.**
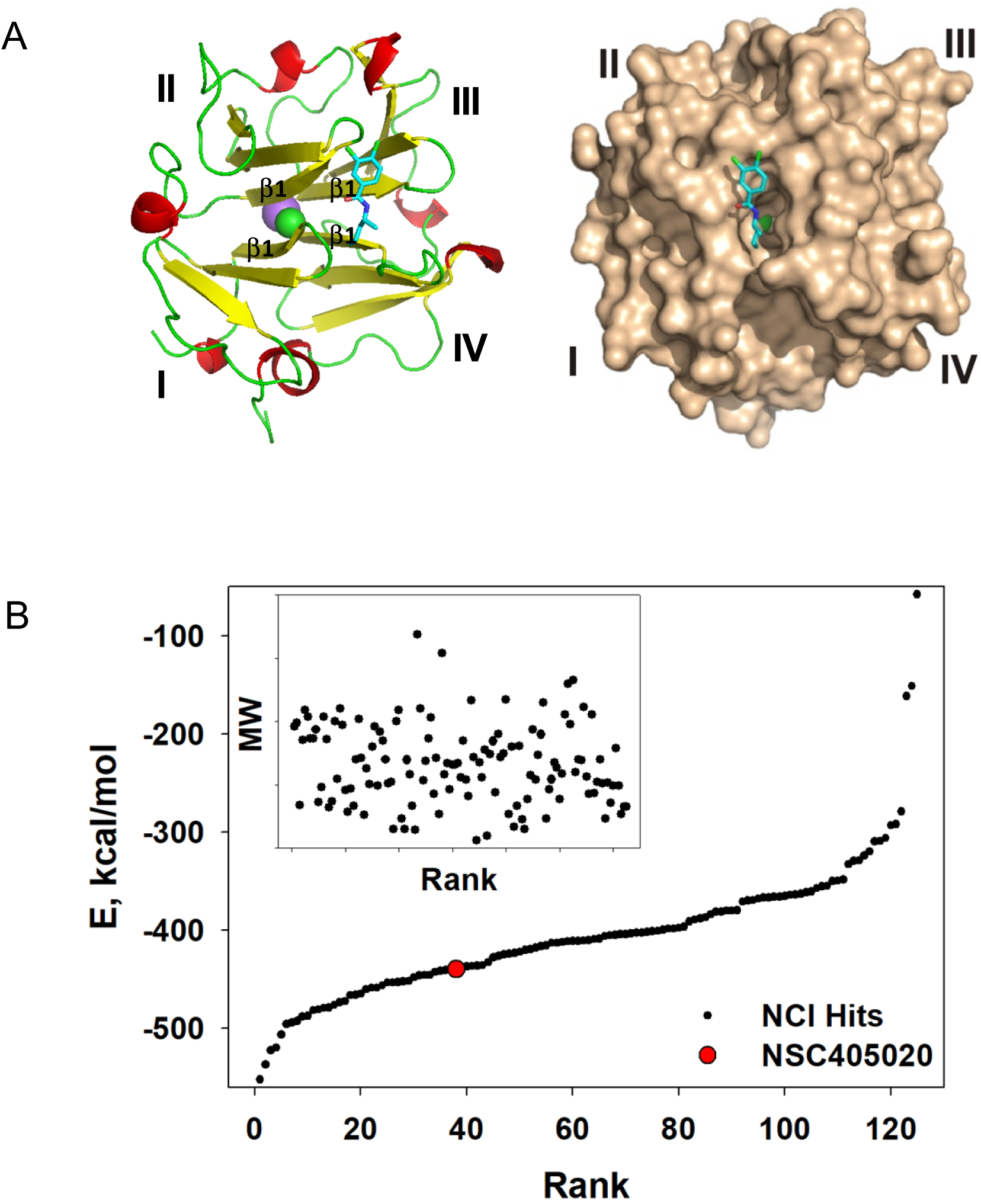
*In Silico* discovery strategy of allosteric modulators of MT1-MMP hemopexin domain. **A.** Putative binding mode of hit compound NSC405020 inside PEX domain (PDB 3C7X) is shown. The docking site was defined in a pocket formed by β-strands 1 (β1) of propeller blades I to IV. **B.** The results of convergent docking (130 hits) of complete NCI DTP ligand database (275,000 compounds) is shown. The ranking of NSC405020 is shown as red circle. **Inset:** hit molecular weight *versus* hit rank distribution.

### Prediction of allosteric ligand binding sites on the surface of protein targets

The Q-MOL *in silico* drug discovery platform has an ultimate advantage over other *in silico* drug discovery methodologies, and this is due to its feature that allows reliable prediction of ligand binding sites on the molecular surface of a protein target (see *Methods*). The Q-MOL ligand binding site prediction was shown to produce relevant predictions in purely theoretical drug discovery research projects ^17, 18^.

The Q-MOL ligand binding sites predictions method can also be applied successfully to so-called "undruggable" targets, e.g. intrinsically disordered proteins such as proto-oncogene cMyc bHLHZ domain (Fig. 7). The structures of known nuclear magnetic resonance (NMR)-validated small molecule cMyc binders ^19^ were successfully used as docking probes to "scan" the molecular surface of the protein target.

**Figure 7.**
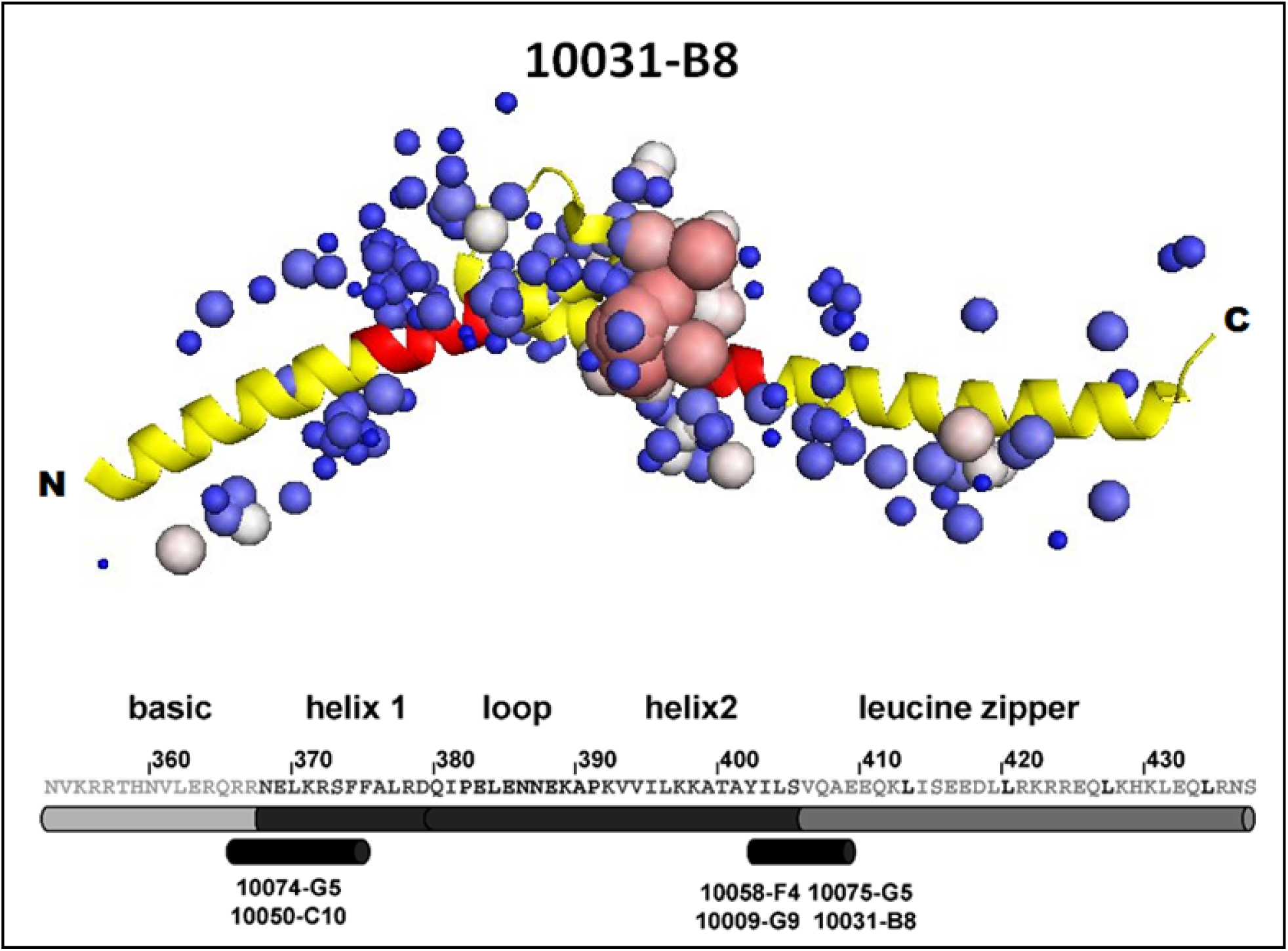
Binding sites mapping of the molecular surface of cMyc protein using *in vitro* validated small molecule ligands as structural probes. Structure of human Myc proto-oncogene protein is shown (PDB 1NKP Myc-Max complex, chain A bHLHZ domain). The Q-MOL ligand binding sites prediction method was applied to Myc protein from Myc-Max complex (Max protein and DNA were removed from simulation). The routine used a small molecule cMyc binder 10031-B8, previously described in ^19^. cMyc molecule is displayed with cartoon style. cMyc amino acid sequence with secondary structure assignments and ligand binding regions assigned using NMR are shown below with corresponding ligand identifiers. The binding regions, corresponding to NMR assignments, are also depicted as red spans on cMyc structure. Probabilities of interactions of 10031-B8 ligands with cMyc amino acid residues are displayed as spheres: large red spheres correspond to high probabilities, small blue spheres correspond to low probabilities.

The Q-MOL ligand binding sites prediction methodology was additionally validated in *de novo* discovery of small molecule modulators of β-catenin ^20^. β-catenin is the key nuclear effector of canonical Wnt signaling pathway, and aberrant Wnt signaling has been closely associated with carcinogenesis ^21^. It has been shown previously that direct targeting of β-catenin armadillo repeat domain is feasible small molecule drug discovery strategy ^22, 23^. A variant of the Q-MOL ligand binding sites prediction routine, where structures of individual 20 amino acids are used as structural molecular probes to scan the molecular surface of a protein target (see *Methods*), was employed to predict locations of potential allosteric binding sites on the surface of β-catenin armadillo repeat domain (Fig. 8). The molecular surface scanning with individual amino acids allows for identification of "hot" spots on the protein molecular surface with excess of stored free energy. This excess surface free energy is one of the major of free energy sources (among solvent effects, surface area changes and others), and it thermodynamically contributes to proteins ability to engage in binding with other proteins and small molecule ligands ^17, 18^. In the case of β-catenin armadillo repeat domain, 3 distinct allosteric sites were detected. The most prominent site (Site C) was selected for protein-ligand docking simulations. The best discovered β-catenin ligand (NSC211416) exhibited direct binding *K_d_* = 55 nM, thermal melting temperature shift *ΔT_m_* = 17°C (Fig. 9), and significant *in vitro* and *in vivo* activity ^20^.

**Figure 8.**
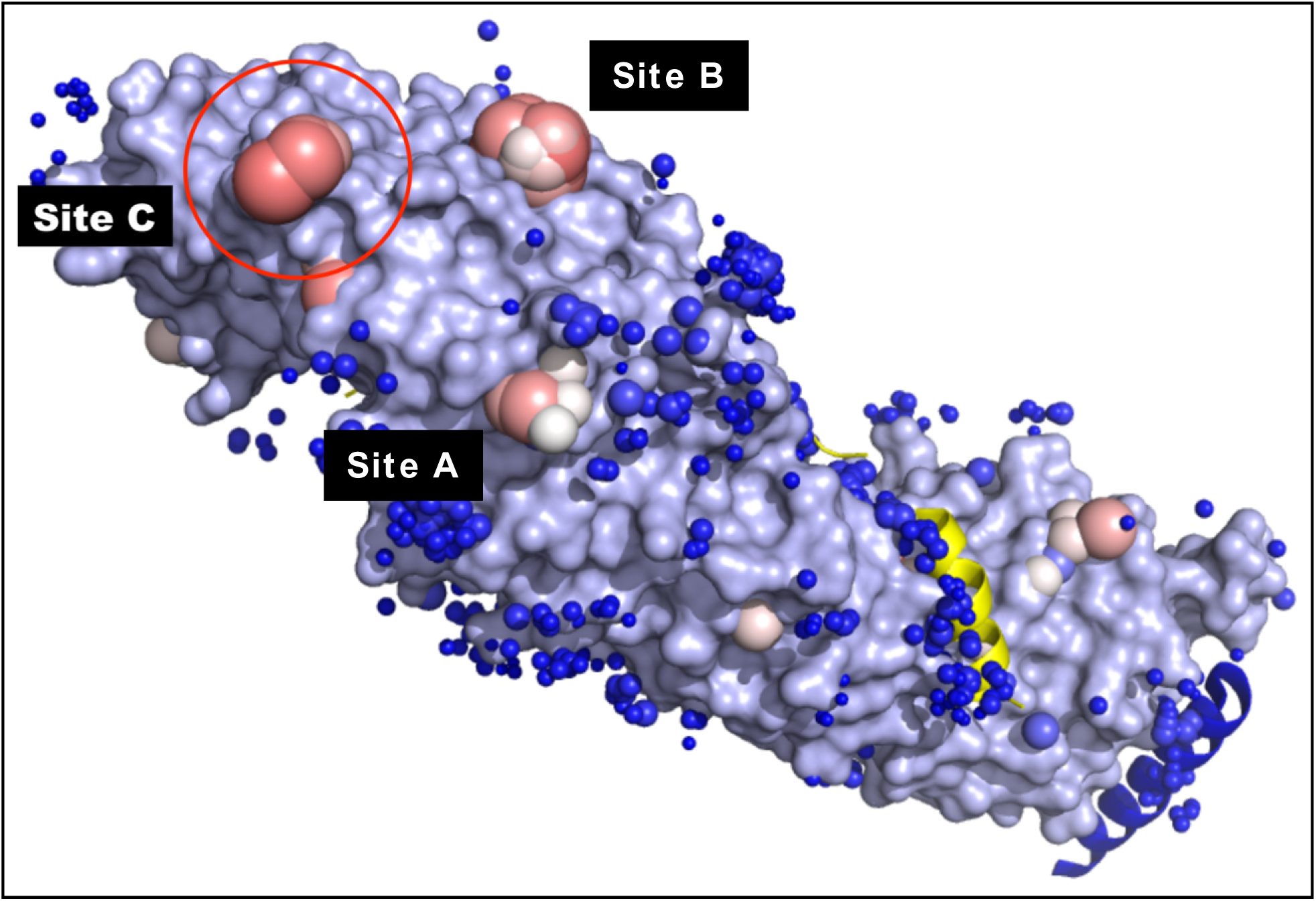
Binding sites mapping of the molecular surface of β-catenin armadillo repeat domain using individual amino acid molecules as structural probes. Structure of human β-catenin armadillo repeat domain in complex with Tcf4 and Bcl9 is shown (PDB 2GL7). β-catenin armadillo repeat is displayed as light blue molecular surface, Tcf4 yellow and Bcl9 blue in cartoon style. Tcf4 and Bcl9 were removed from the complex before application of Q-MOL MSBM routine. Probabilities of interactions of individual amino acid molecules with the surface of protein target are displayed as spheres: large red spheres correspond to high probabilities, small blue spheres correspond to low probabilities. Predicted allosteric binding sites (clusters of high probability values) are labeled as sites A, B, C. Site C, selected for protein-ligand docking simulation is highlighted with red circle.

**Figure 9.**
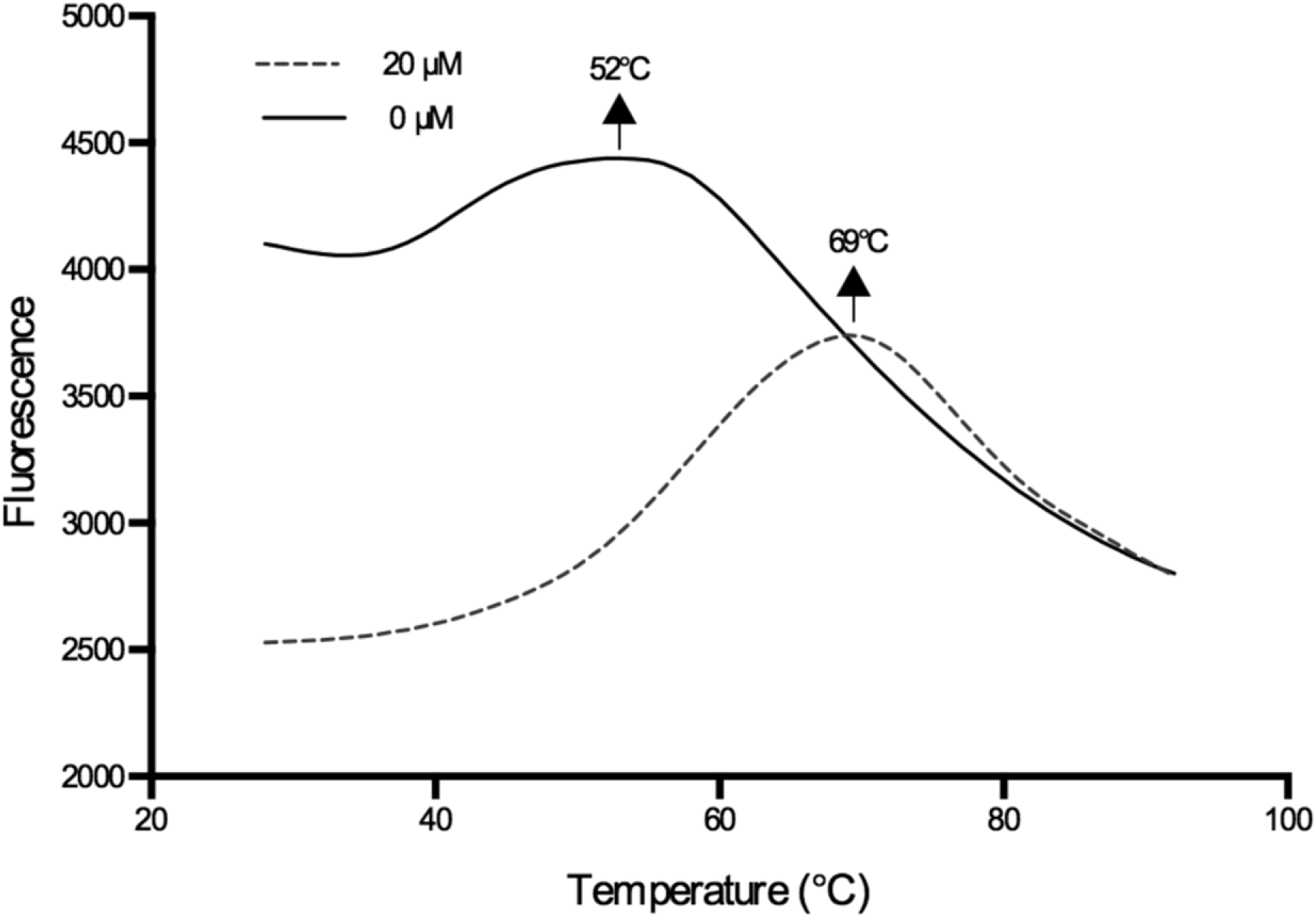
Thermal melting profile of recombinant wild type human β-catenin in complex with NSC211416. 2.5 μM purifed recombinant wild-type human β-catenin protein, 20 μM NSC211416.

**Figure 10.**
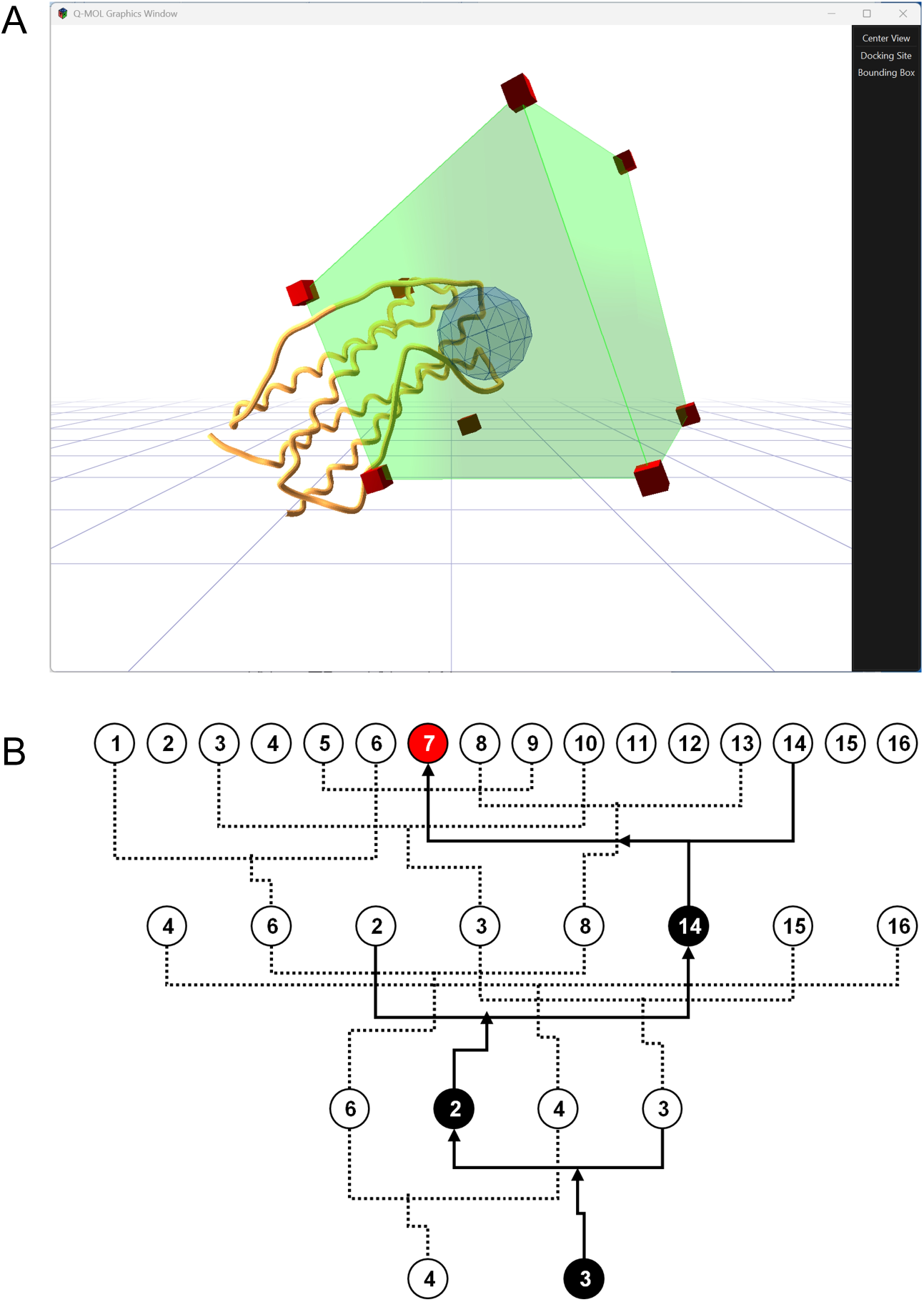
Q-MOL protein-ligand docking setup and compound library traversal logic. A. Ligand docking configuration setup for Q-MOL docking project. Receptor (IL-11, PDB 6O4O) is shown in cartoon style. The starting point of ligand docking simulation (the docking site) is shown is purple wired sphere, boundary condition is displayed as green box. **B. Schematic diagram of application of chemical BSP to virtual ligand screening.** The original compound library (16 compounds) is clustered by chemical similarity measure into a layered tree-like data structure – binary space partitioning tree. The VLS is initiated at the lowest level of the tree, containing only 2 compounds. The tree branches of the best scoring compound (#3) are then traversed toward the top of the tree. At each branching point, the leaf containing the best scoring compound is selected: 3⇨2⇨14⇨7. The total number of docked ligands is 5 (2+3), the best scoring ligand is #7. The diagram demonstrates the idea, not the exact implementation.

### Targeting IDPs and highly-flexible proteins

One of the lucrative aspects of *in silico* drug discovery is to approach protein targets which are notoriously difficult to work with *in vitro*, or which are operating within cellular systems that are not possible to efficiently scale and manipulate in a laboratory (e.g. growing primary neurons in culture). These protein targets are often highly flexible or IDPs that belong to protein families of transcription factors, protein adaptors, flexible regulatory domains and interfaces of large enzymes (e.g. PEX domain of MT1-MMP), and many others. The reality is that intrinsically disordered and highly flexible proteins constitute the bulk of important drug targets. Only a few of these protein targets with known structure contain canonical druggable binding pockets. The rest of the bulk are considered undruggable, though it is understood that many of them may contain "cryptic" binding sites that are realized in specific protein conformations ^24^. There exist computational methodologies that attempt to detect such sites with varying degree of success (reviewed in ^1^). However, after a putative binding site is identified, the next step presents another problem: how to correctly account for specific protein conformational changes that result in the predicted binding pocket during protein-ligand docking simulations.

The direct structure modeling methods (e.g. molecular dynamics) are currently impractical for any kind of practical applications because they require immense computational resources and critically lack information about details of specific structural features during the conformational transaction of a given protein target. Those details include effects of cell-type dependent environment, presence of phospholipid membrane and/or other binding partners, solvent, and a "crowding effect" ^25, 26^. As a more computationally practical alternative, the machine learning (ML) methods are used. They work by transforming an input vector or a matrix of structural parameters into an output of similar kind. A transform (e.g., a neuronal network), which is applied to input parameters, is essentially a black box that has been trained on a limited set of experimental data and has no connection to physics and thermodynamics of a real system. As a result, this approach fails when a protein target of interest does not belong to any of structural families of proteins that were used for the transform training.

The approaches, briefly discussed above, are the two extreme ends of the methodology. What if a good [enough] solution can be found in-between? If protein flexibility cannot be reliably modeled explicitly, it could be modeled implicitly with reasonable simplifications to the system. In contrast to black box ML methods, implicit protein flexibility modeling needs to rely explicitly on force field parameters and structural features of a target protein. To this end, Q-MOL protein-ligand docking methodology has employed the energy landscape theory of protein folding (ELT) ^2, 27–35^.

Briefly, the ELT is a statistical description of a protein’s potential energy surface. It assumes that folding occurs through organizing ensembles of structures rather than through only a few uniquely defined structural intermediates. The ELT describes the process of protein folding into its potentially many [local] energy minima states as a *reversible* journey through a complex energy landscape. This landscape can be visualized theoretically as a funnel-like shape (the folding funnel). Experimentally, we can obtain a cumulative "cross section" of the folding funnel by a thermal melt assay where the funnel is "projected" on a plane *assay readout* vs *T_M_*, where the *assay readout* is some assay output parameter (e.g. fluorescence), which is proportional to the number of specific protein conformations at given melting temperature *T_M_*(e.g. Fig. 9). The overall folding path is guided by the amino acid sequence, interactions within the molecule, and importantly, the local environment such as the presence of ligands, other binding proteins and the crowding effect. The dominant conformation of the most populated ensemble represents an apparent native state. This conformational ensemble is in dynamic equilibrium with other ensembles, and depending on a local environment, it can rapidly shift to a "new" apparent native state. Therefore, a solved (e.g. X-ray or NMR) protein structure is apparently just a singular "snapshot" from a vast array of conformational states existing in a dynamic equilibrium with each other. Thus, the "native" structure of a protein is a metastable structure that exists only within a particular cellular context or an experimental environment. The metastable nature of a native state of a protein leads to the lack of critical structural data that are needed to train an AI/ML model or a scoring function. As a result, *in silico* methods directly employing AI/ML and scoring functions will fail to deliver correct results for protein targets either missing from a training set, or missing specific structural contexts within a set.

Though the theoretical statements of ELT are clearly defined and are confirmed in many experimental studies, ELT does not provide exact instructions how its statements should be implemented computationally. One important conclusion that can be drawn from ELT about the protein-ligand binding process is that it is ligand-centric. A ligand binds to the best fit conformation available from protein folding funnel as in a lock-and-key model. If binding is strong enough and the ligand is available in sufficient quantities the protein conformation in complex with a ligand becomes the dominant entity. The change in protein function as a result of the binding event is then reflected in a readout of a biochemical or cell-based assay. Importantly, the ligand’s chemical structure encodes structural information about specific interactions with a unique receptor conformation even if the exact binding mode information is not available.

The Q-MOL protein-ligand docking methodology computationally mimics this ligand-centric nature of the ligand binding. The protein-ligand docking simulation runs in the internal coordinate space of a ligand, however its computed binding energy with a receptor (a known "snapshot" structure) is represented as a multidimensional parametric hyperspace. The simulation initially starts with a single energy "dimension", and as the simulation progresses and new protein-ligand interactions are discovered, new dimensions of binding energy are added to the multidimensional parametric receptor representation. This collection of binding energy dimensions implicitly (and presumably) represents all possible conformations on a protein potential energy surface that are amenable for binding of a given ligand. The resulting computed binding energy of an individual ligand is used directly to prioritize ligands. Q-MOL protein-ligand docking does not employ a trained ligand scoring function for ranking of hits. Insets in figures 2, 3 and 6 are shown to demonstrate that the resulting hit ranking is not affected by a ligand size. In contrast, many classic hit scoring functions are using ligand size penalty terms to avoid rank skewing with the size of a ligand. The exact details of the implementation of the algorithm will be described elsewhere.

The ELT-based implicit receptor flexibility treatment provides several unique capabilities to the Q-MOL *in silico* drug discovery platform. The most important and unique capability is the ability to successfully dock ligands into putative and cryptic binding sites lacking well-defined druggable pocket features. In fact, the Q-MOL protein-ligand docking can be successfully applied against completely flat and structurally featureless surface sites ^9, 20^ (Fig. 1, Fig. 8).

Yet another related unique feature is the ability to reliably predict allosteric and cryptic binding sites ^17, 20^ (Fig. 7, Fig. 8). The binding sites prediction is absolutely critical for *in silico* drug discovery because highly flexible and intrinsically disordered protein targets can use distinct surface binding interfaces, depending on cellular context. This information is not readily available *a priori* or not easy to obtain via conventional biochemical and biological methods (e.g. site-directed mutagenesis). This sites prediction feature is based on the protein-ligand docking with ETL-based protein flexibility treatment, which is systematically applied across the entire molecular surface of a receptor (see Methods). The Q-MOL site detection protocol "scans" the molecular surface of target protein using a ligand molecule as structural probe. Essentially, the docking of the probe detects excess (if any) free protein surface energy. This surface energy is normally used by a protein to drive thermodynamically favorable binding. The feature allows binding sites detection not only when a specific ligand is known (e.g., Fig. 7), but also in the absence of such information. In the latter case, structures of 20 individual amino acids are used as docking probes (Fig. 8, Fig. 11).

**Figure 11.**
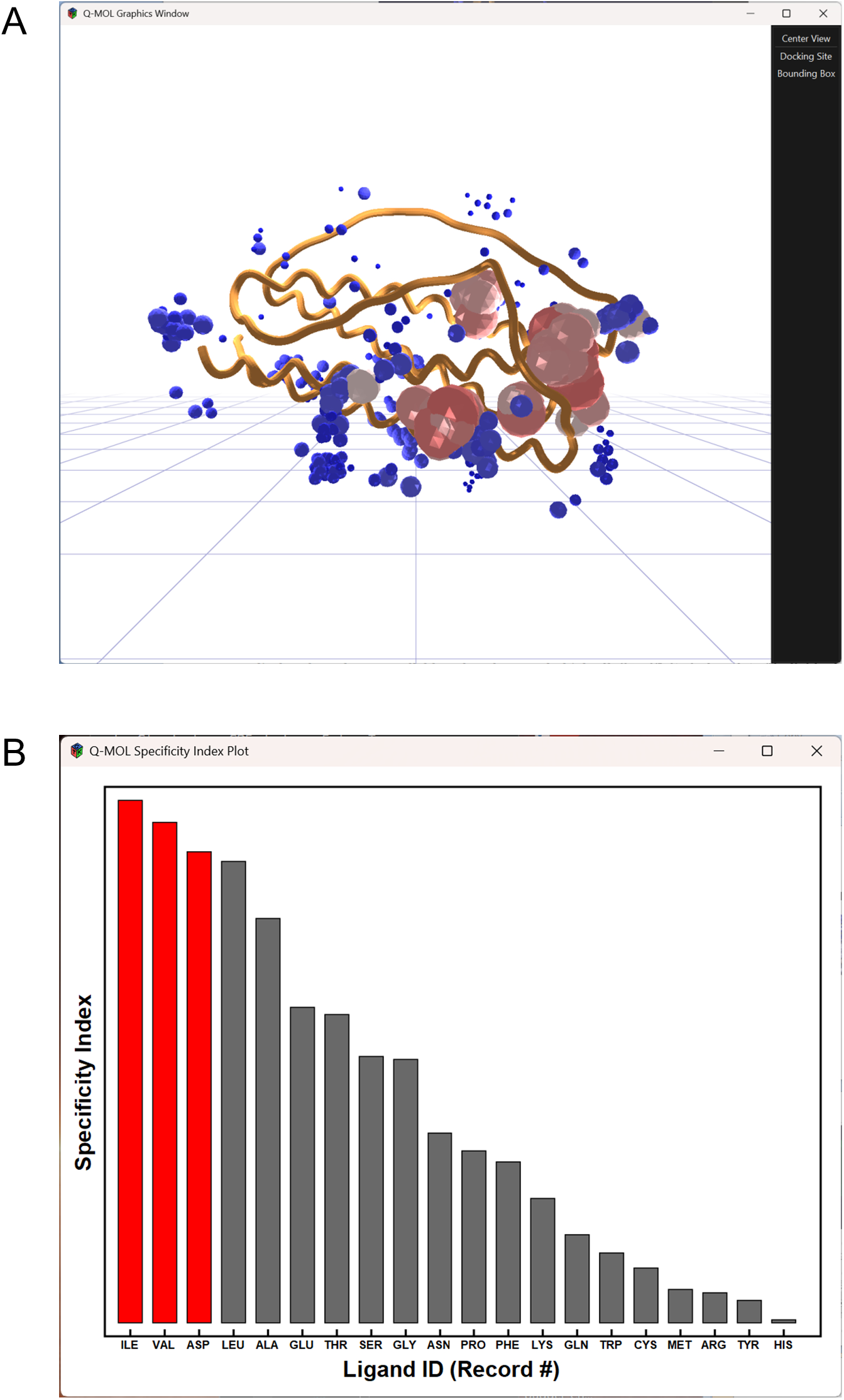
Detection of allosteric binding sites. A. Visualization of molecular surface scan results. Receptor (IL-11, PDB 6O4O) is shown in cartoon style. Probabilities of binding of individual amino acids (Ile, Val, Asp, see panel **B**) at specific sites on the molecular surface are displayed as spheres: small, blue - low, large, red - high. **B. Specificity index plot of individual amino acids used as scanning structural probes.** Specificity index was computed for each amino acid, amino acids were sorted by specificity index (see *Methods* for details).

Because of the nature of the Q-MOL protein-ligand docking methodology, *de novo* drug discovery workflow with Q-MOL software follows specific sequence of steps. For a new protein target, putative binding sites are detected first by the described molecular surface scanning using either a known ligand (e.g. obtained from a functional assay) or 20 individual amino acids as docking probes (e.g. Fig. 7, Fig. 8, Fig. 11). After selection of a docking site, the next computational step is the primary VLS. The primary VLS identifies potential binders toward a selected binding site for all possible receptor conformations in proximity to that binding site (e.g. Fig. 2, Fig.3, Fig. 5). As a result, the primary VLS docking curve represents a collection of ligands with the best (lowest) binding free energies computed from the ELT-based representation of a receptor as a multidimensional potential energy landscape. Accordingly, each ligand from the docking curve is a potential binder, though a lower relative binding free energy indicates a stronger binding possibility (but not necessarily better selectivity). Thus, the ligands of interest across all of the docking curve could then be selected for biological validation.

If a protein target is known to perform alternative functions or have alternative binding partners in different cell types, then the selected ligands need to be assayed in each specific cell type. The biological validation in different cell types will likely produce different hit sets (Fig. 12). Importantly, Q-MOL VLS can produce hits that can behave as either activators or inhibitors in a functional or biochemical assay. This information cannot be deduced from a docking simulation because of the implicit treatment of target flexibility.

**Figure 12.**
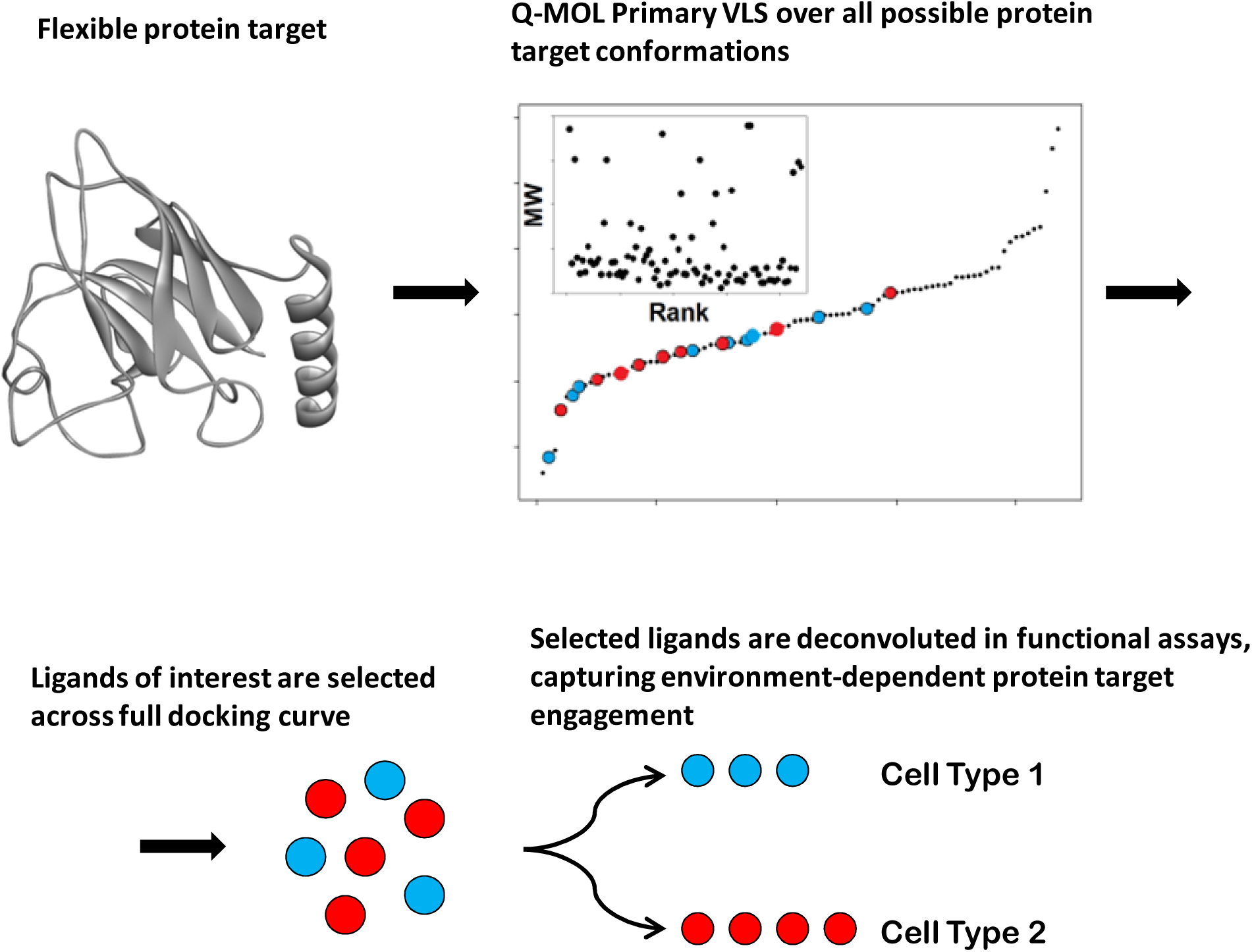
Schematic diagram of functional deconvolution of Q-MOL primary VLS docking hits against a flexible protein target that is conformationally dependent on cellular environment. After Q-MOL primary VLS is applied to a protein target, predicted hits of interest are selected across full docking curve. The selected hits form the pool of ligands that at are predicted to bind to one or more conformations from potentially many conformational ensembles that represent folding energy landscape of a given flexible receptor. Different cellular environments promote specific conformational ensembles of the receptor via interactions with its cell-type specific binding partners (cellular context). A specific functional cell-based assay for each cell type detects modulation of function when selected ligands specifically interact with context-dependent conformations of a receptor.

The set of biologically validated hits from primary VLS represents ligands that bind unique receptor conformations from an ensemble of conformations that are biased toward some local energy minimum (environment-dependent apparent native state). However, these ligands might be interacting with receptor conformations relatively distant from that of the apparent native state, and as a result, they do not modulate efficiently desired function of a protein target. There could be several reasons for this including lack of chemotypes in the library used for VLS, and, importantly, the conformational distance of the "snapshot" receptor structure from the apparent native state. Thus, the structural space of this conformational ensemble still needs to be fully sampled with additional ligands. To this end, several hundreds of closest chemical analogues of a functionally validated hit are searched in a database to form a focused library. The focused library is then docked and new hits are selected for biological validation (see Methods) ^12^ (Fig. 4). Because the focused library is sampling a single conformational ensemble, the top (lowest energy) binders are selected for assay. This is different from hits selection logic for a primary VLS when hits are selected along the complete docking curve.

### Applicability of Q-MOL Allosteric Sites Prediction and Ligand Docking Methodologies to Polynucleotide Molecules

To validate the universal nature of ELT implementation and reparameterized ligand docking OPLS force field, the protein-optimized computational protocols were applied "As Is" to non-protein polymeric molecules such as ribonucleic acids (RNA). To this end, two non-coding viral RNA molecules of known structure were investigated: human immunodeficiency virus (HIV) - 1 core packaging signal (PDB 2N1Q, NMR)^36^ and exonuclease resistant RNA from Zika virus (PDB 5TPY, X-ray structure)^37^. The HIV-1 core packaging signal is a cis-acting RNA element located at the 5’ leader end of the viral genome. It directs the packaging of unspliced viral RNA into assembling virus particles by interacting with the Gag protein^38^. The HIV-1 core packaging signal is highly conserved and represents an amenable target for potential therapeutic intervention aiming to block virus assembly^39^. Zika virus (ZIKV) produces exonuclease-resistant RNAs (xrRNAs) in its 3’ untranslated region (UTR) that act as mechanical roadblocks against host cell 5’-3’ exoribonucleases (specifically Xrn1)^37^. These structures, specifically xrRNA1 and xrRNA2, utilize a complex, highly stable fold with a molecular "ring" to halt degradation, producing subgenomic flavivirus RNAs (sfRNAs)^40^. sfRNAs are essential for viral pathogenicity and evasion of the host immune system. The resistance of these sfRNAs to host cell exoribonucleases results in their accumulation. It has been shown that disrupting these specific RNA structures via mutations can abolish their accumulation, significantly impacting the virus’s ability to infect. Thus, a therapeutic intervention aimed to interfere with flavivirus sfRNAs stability might prove to be a highly effective and precise strategy for treating this viral infection.

The standard Q-MOL drug discovery protocols were applied to both RNA structures. The probable allosteric binding sites were predicted by applying molecular surface scanning protocol using structures of individual amino acids as docking probes (see *Methods*). In both cases, the simulation produced statistically strong signal identifying specific areas on the surface of the molecules (Figure 13A, Figure 14A).

**Figure 13.**
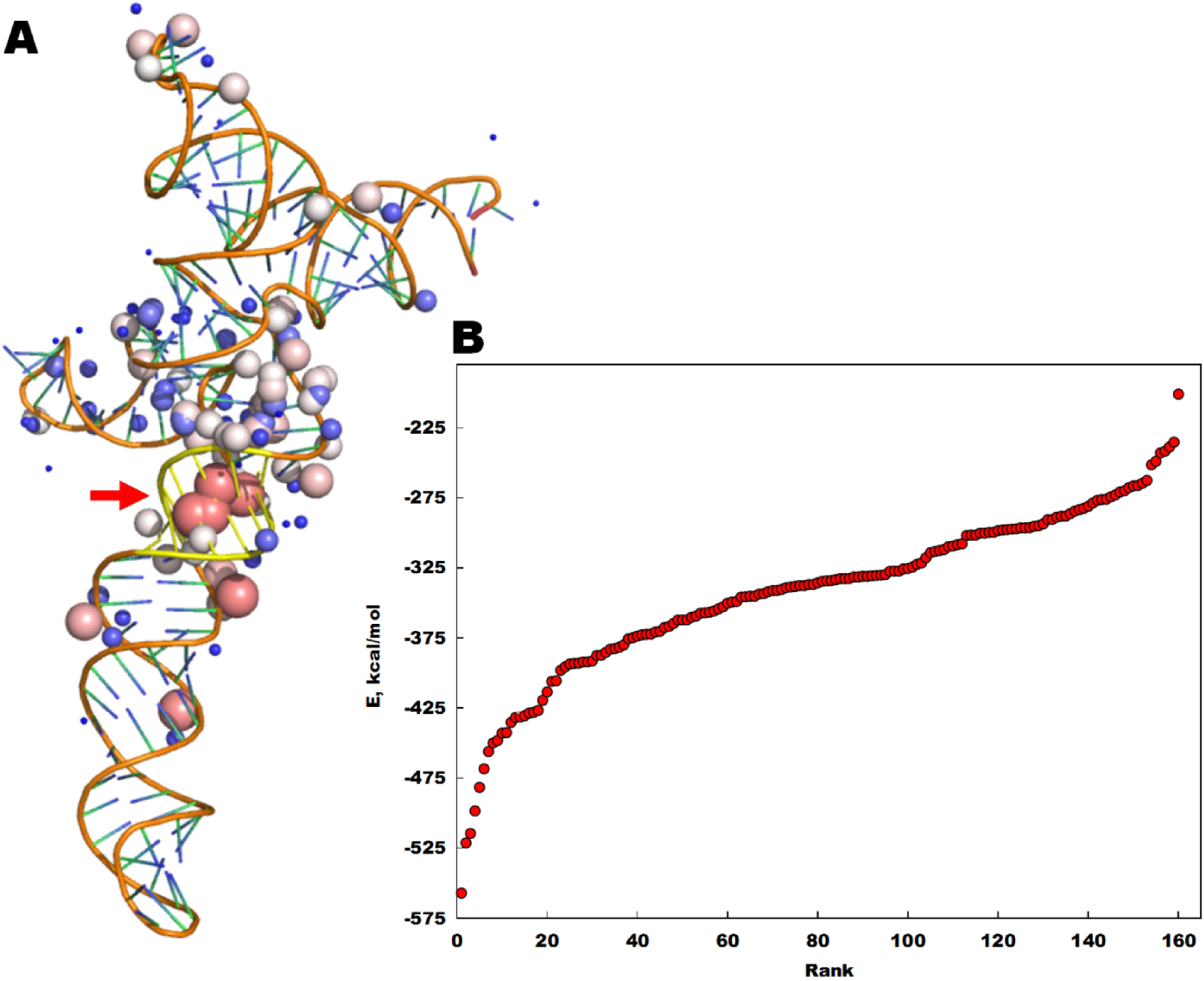
Predicted small molecules binding site and primary VLS results against HIV-1 Core Packaging Signal RNA. A. NMR structure of HIV-1 Core Packaging Signal RNA (PDB 2N1Q). The putative binding site was predicted by Q-MOL molecular surface scanning using individual amino acid structures as docking probes (see *Methods*). Probabilities of binding across the molecular surface are shown as spheres (red/large – high probability, blue/small – low probability). The nucleotides, forming predicted allosteric site (cluster of high probabilities, indicated with a red arrow), are highlighted yellow. The following nucleotide stretches define the predicted binding site: U(230)CUCGAC(236) … G(282)GCGAC(287). The residue numbering corresponds to that of PDB 2N1Q. **B. Primary Q-MOL VLS results.** The complete NCI DTP SDF library was used as a source of ligands (≈ 275,000 structures). The primary Q-MOL VLS has converged to 162 individual hits. The ligand docking curve is shown. The docking site was defined around the highlighted yellow residues (indicated with a red arrow). *E*, relative binding energy, kcal/mol; *Rank*, ligand ranking based on the sorting by relative binding energy.

**Figure 14.**
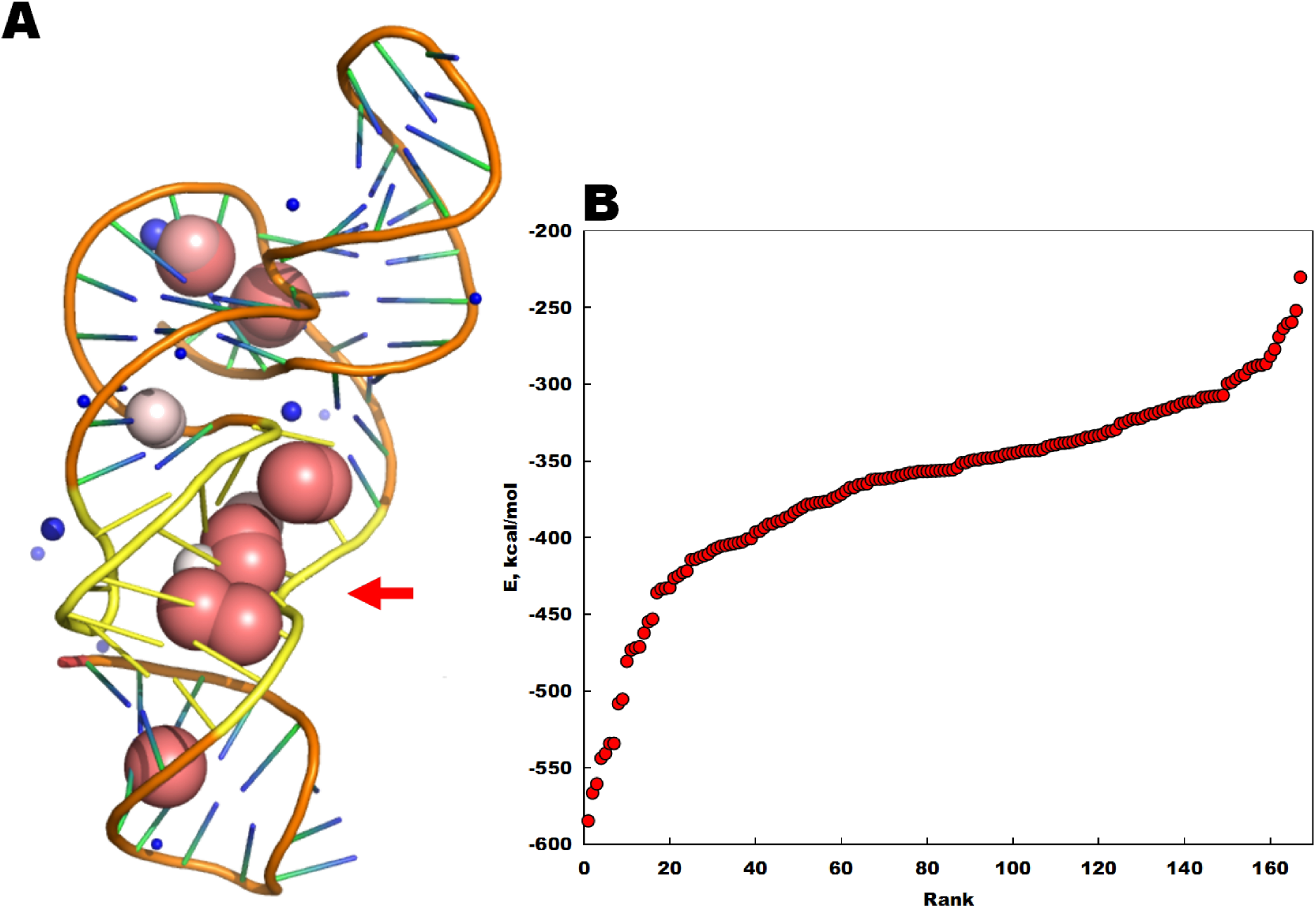
Predicted small molecules binding site and primary Q-MOL VLS results against an exonuclease resistant RNA from Zika virus. A. X-ray structure of exonuclease resistant RNA from Zika virus (PDB 5TPY). The putative binding site was predicted by Q-MOL molecular surface scanning using individual amino acid structures as docking probes (see *Methods*). Probabilities of binding across the molecular surface are shown as spheres (red/large – high probability, blue/small – low probability). The nucleotides, forming predicted allosteric site (cluster of high probabilities, indicated with red arrow), are highlighted yellow. The following nucleotide stretches define the predicted binding site: U(29)UGGGGAAA(37) … A(52)ACCCC(57). The residue numbering corresponds to that of PDB 5TPY. **B. Primary Q-MOL VLS results.** The complete NCI DTP SDF library was used as a source of ligands (≈ 275,000 structures). The primary Q-MOL VLS has converged to 169 individual hits. The ligand docking curve is shown. The docking site was defined around the highlighted yellow residues (indicated with a red arrow). *E*, relative binding energy, kcal/mol; *Rank*, ligand ranking based on the sorting by relative binding energy.

For the HIV-1 core packaging signal, the allosteric binding site is predicted to be formed by the following nucleotide stretches: U(230)CUCGAC(236) … G(282)GCGAC(287). The residue numbering corresponds to that of PDB 2N1Q.

In the case of Zika virus exonuclease resistant RNA, the allosteric binding site is predicted to be formed by the following nucleotide stretches: U(29)UGGGGAAA(37) … A(52)ACCCC(57). The residue numbering corresponds to that of PDB 5TPY. The primary Q-MOL VLS was performed using the complete NCI DTP SDF library (≈ 275,000 compounds) (Figure 13B, Figure 14B). The docking sites for RNA-ligand docking simulations were defined around the predicted allosteric binding sites. The predicted hit structures (162 hits, HIV-1 core packaging signal; 169 hits, Zika virus exonuclease resistant RNA) were bucket-clustered by similarity, and the representative ligands of the most populated cluster buckets were annotated when possible (Table 1, Table 2).

**Table 1.**
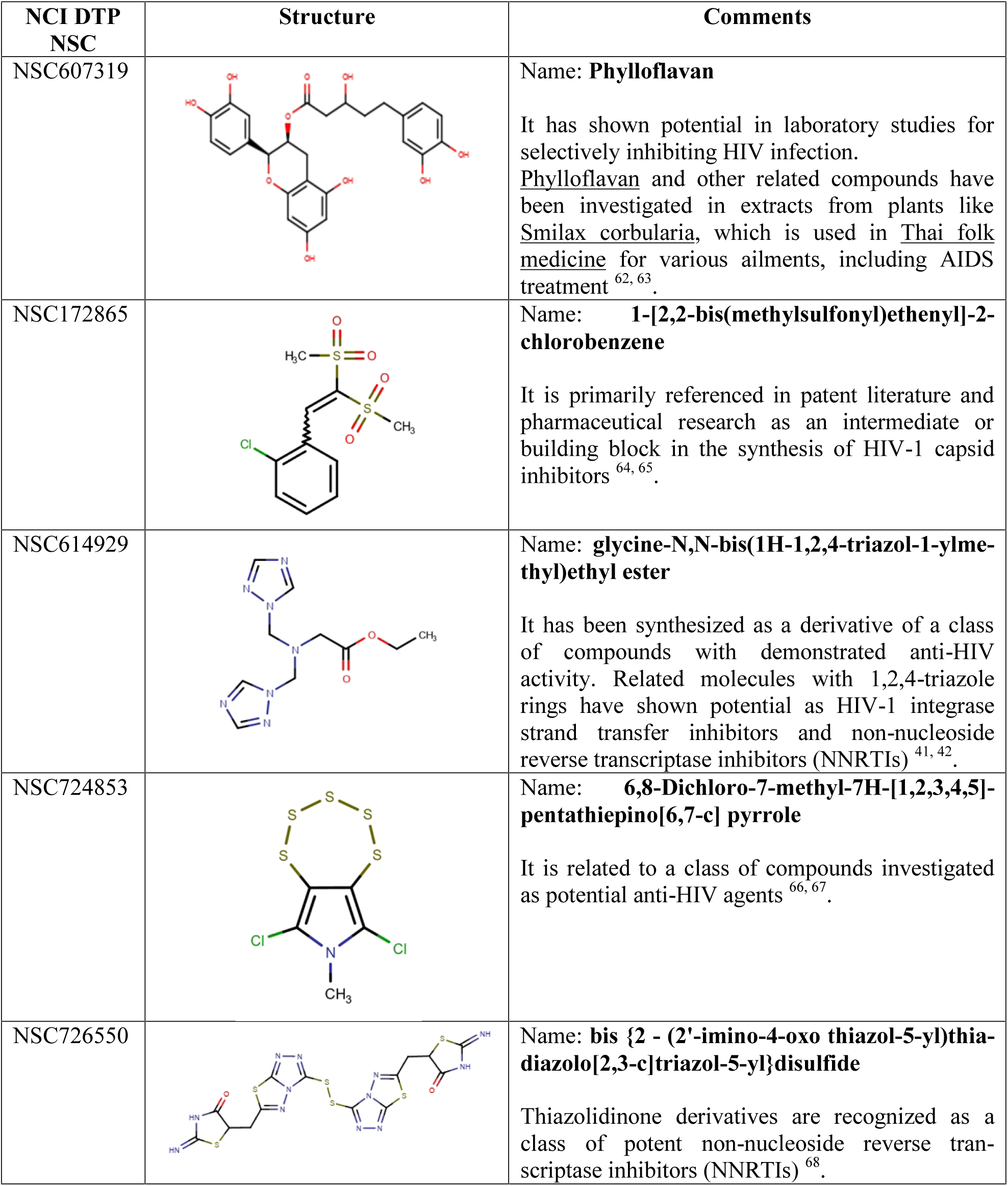
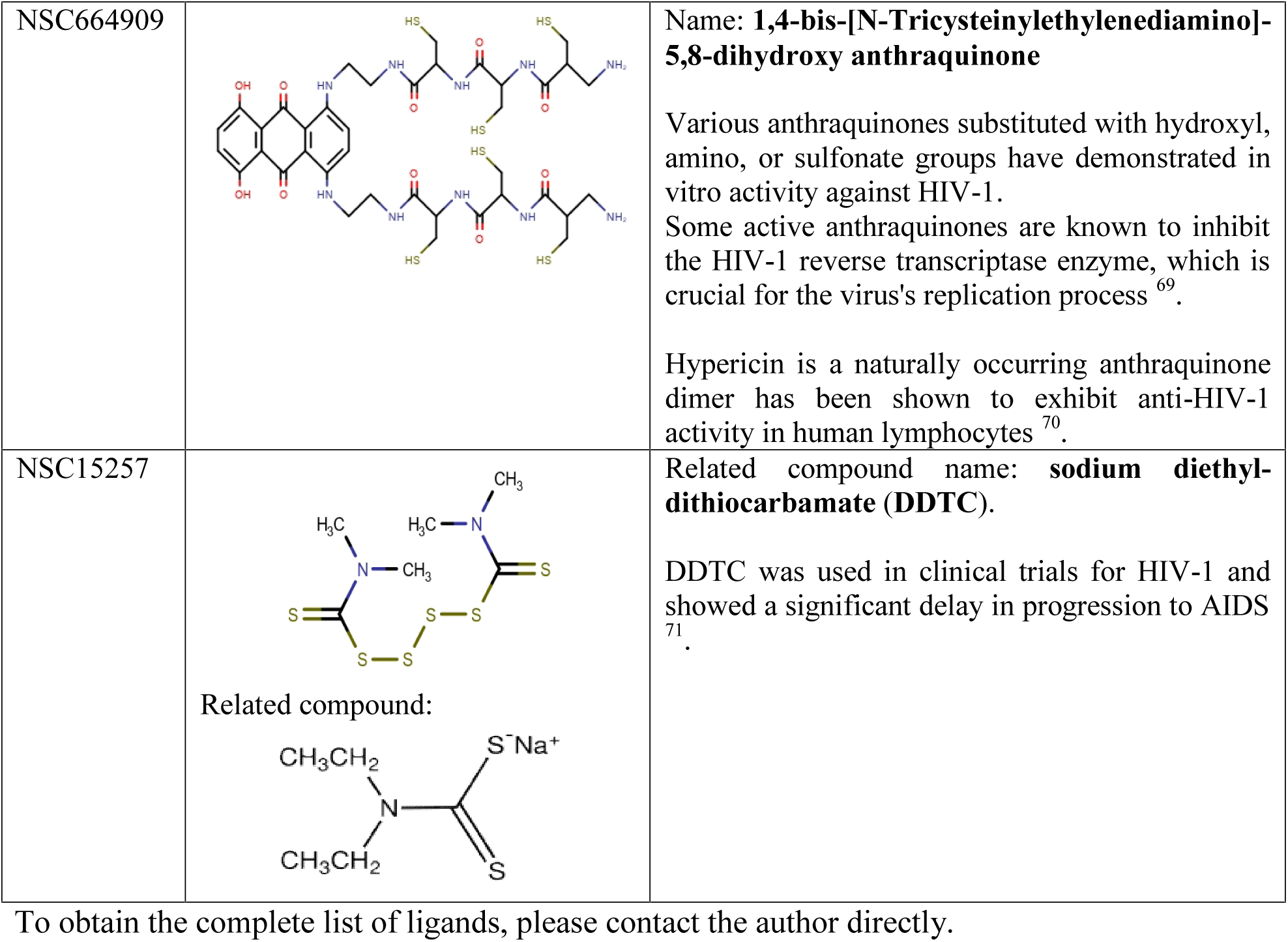
Selected annotated ligands from NCI DTP library identified as hits by Q-MOL primary VLS against HIV-1 Core Packaging Signal RNA.

**Table 2.**
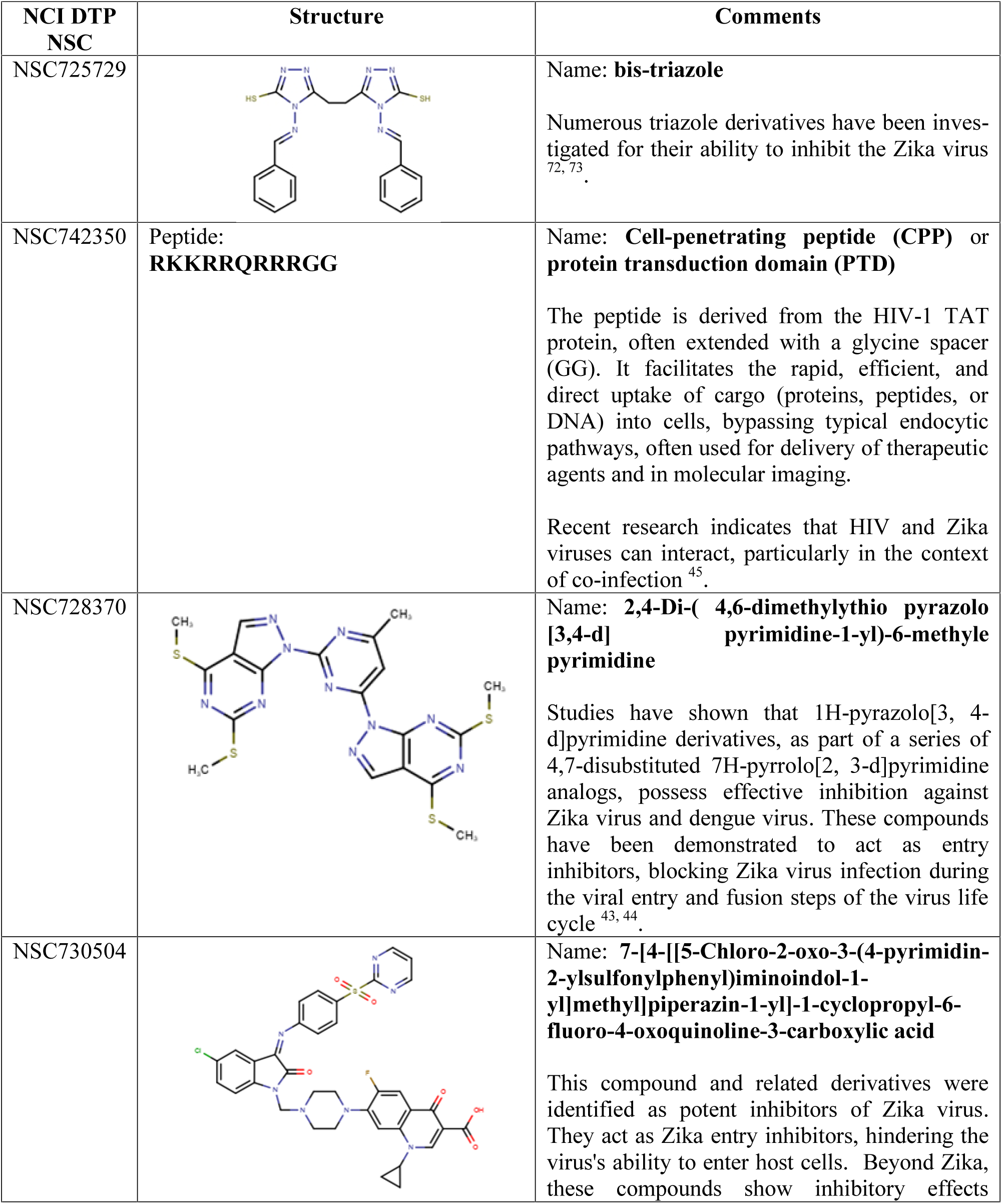
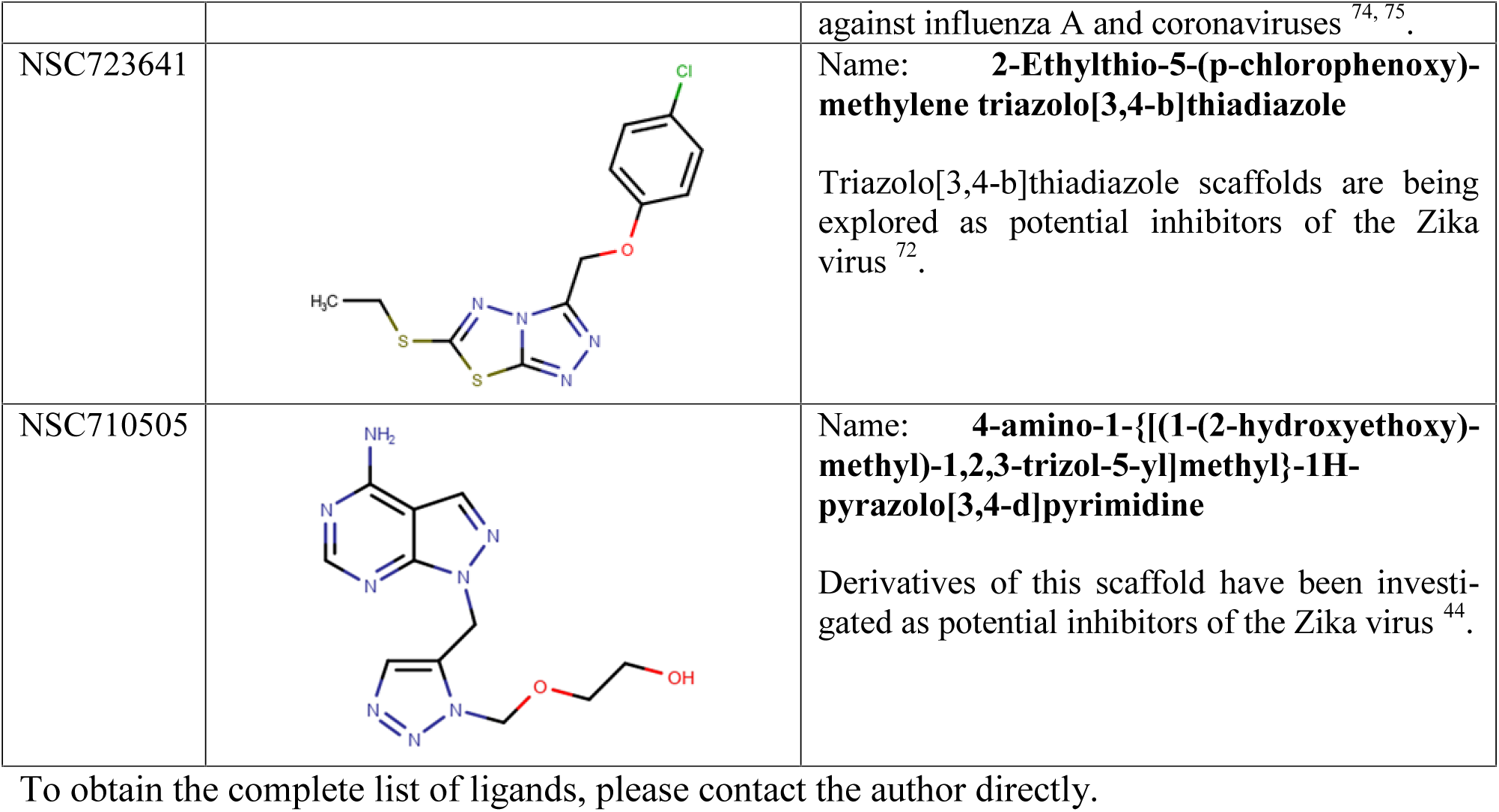
Selected annotated ligands from NCI DTP library identified as hits in Q-MOL primary VLS against exonuclease resistant RNA from Zika virus.

| NCI DTP NSC | Structure | Comments |
| --- | --- | --- |
| NSC725729 |  | Name: **bis-triazole**<br>Numerous triazole derivatives have been investigated for their ability to inhibit the Zika virus <sup>72, 73</sup>. |
| NSC742350 | Peptide:<br>**RKKRRQRRRGG** | Name: **Cell-penetrating peptide (CPP)** or **protein transduction domain (PTD)**<br>The peptide is derived from the HIV-1 TAT protein, often extended with a glycine spacer (GG). It facilitates the rapid, efficient, and direct uptake of cargo (proteins, peptides, or DNA) into cells, bypassing typical endocytic pathways, often used for delivery of therapeutic agents and in molecular imaging.<br>Recent research indicates that HIV and Zika viruses can interact, particularly in the context of co-infection <sup>45</sup>. |
| NSC728370 |  | Name: **2,4-Di-( 4,6-dimethylthio pyrazolo [3,4-d] pyrimidine-1-yl)-6-methyle pyrimidine**<br>Studies have shown that 1H-pyrazolo[3, 4-d]pyrimidine derivatives, as part of a series of 4,7-disubstituted 7H-pyrrolo[2, 3-d]pyrimidine analogs, possess effective inhibition against Zika virus and dengue virus. These compounds have been demonstrated to act as entry inhibitors, blocking Zika virus infection during the viral entry and fusion steps of the virus life cycle <sup>43, 44</sup>. |
| NSC730504 |  | Name: **7-[4-[[5-Chloro-2-oxo-3-(4-pyrimidin-2-ylsulfonylphenyl)iminoindol-1-yl]methyl]piperazin-1-yl]-1-cyclopropyl-6-fluoro-4-oxoquinoline-3-carboxylic acid**<br>This compound and related derivatives were identified as potent inhibitors of Zika virus. They act as Zika entry inhibitors, hindering the virus's ability to enter host cells. Beyond Zika, these compounds show inhibitory effects |
|  |  | against influenza A and coronaviruses <sup>74, 75</sup>. |
| NSC723641 |  | Name: **2-Ethylthio-5-(p-chlorophenoxy)-methylene triazolo[3,4-b]thiadiazole**<br>Triazolo[3,4-b]thiadiazole scaffolds are being explored as potential inhibitors of the Zika virus <sup>72</sup>. |
| NSC710505 |  | Name: **4-amino-1-{[(1-(2-hydroxyethoxy)-methyl)-1,2,3-triazol-5-yl]methyl}-1H-pyrazolo[3,4-d]pyrimidine**<br>Derivatives of this scaffold have been investigated as potential inhibitors of the Zika virus <sup>44</sup>. |

To obtain the complete list of ligands, please contact the author directly.

These RNA-ligand docking experiments are obviously lacking *in vitro* validation, however the limited annotated results demonstrate an excellent correlation between annotated small molecule hits and the viral origins of RNA structures. Moreover, the RNA-ligand docking identified hits that were only relatively recently characterizes as having potent anti-HIV and anti-Zika antiviral properties. For instance, NSC614929 (Table 1) is a derivative of compounds containing 1,2,4-triazole rings. The compounds, containing 1,2,4-triazole rings, have shown potential as HIV-1 integrase strand transfer inhibitors and non-nucleoside reverse transcriptase inhibitors^41, 42^. The mechanism of action of these compounds might need to be re-visited.

In case of Zika virus (Table 2), the analogues of compound NSC728370 have been shown to possess effective inhibition against Zika virus^43, 44^. Of interest, the cell-penetrating peptide (CPP) (NSC742350, Table 2), derived from HIV-1 TAT protein, has been identified as another binder of Zika exonuclease resistant RNA. This finding might be of importance for the understanding of Zika virus interactions with host immune systems as recent research indicates that HIV and Zika viruses can interact, particularly in the context of co-infection^45^.

### Application of Q-MOL Allosteric Sites Prediction and Ligand Docking Methodologies to "undruggable" IDP target c-Myc

The prediction of binding sites, using structures of known NMR-confirmed ligands as docking probes (Fig. 7, see *Methods*), was further validated with more general approach using structures of individual amino acids as docking probes for the molecular surface scanning (Fig. 15A, see *Methods*). In addition to re-confirmation of ligand binding site detected by 10031-B8 ligand (Fig. 7; Fig. 15A, docking site definition), the amino acid scan has also identified additional putative binding site on Myc-Max interface. For the docking experiments, the docking site, identified by both 10031-B8 and amino acids scanning, was chosen (Fig. 15A). The Human Metabolites library was used as a ligand source^46^.

**Figure 15.**
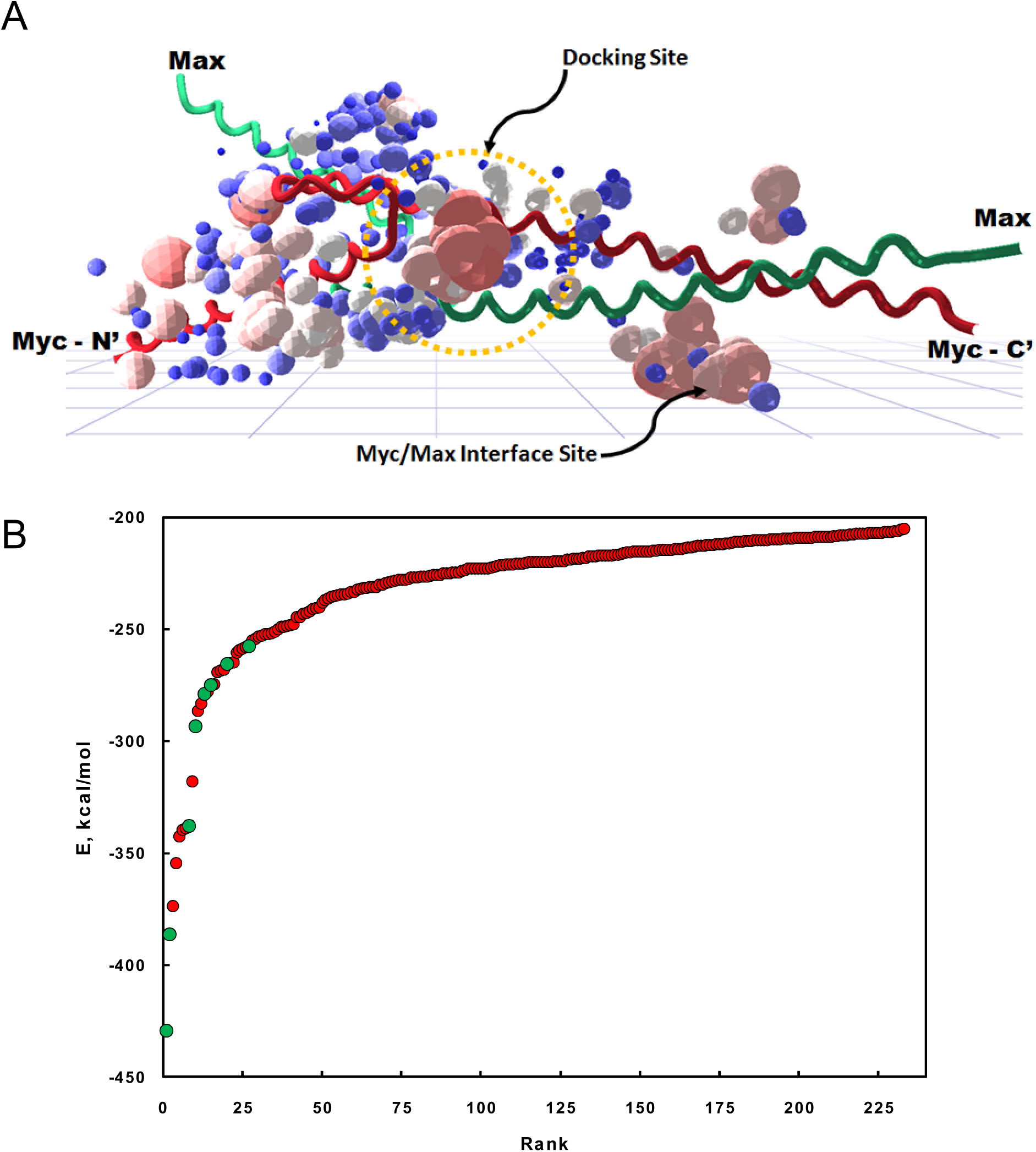
Predicted allosteric sites and human metabolites library VLS results against c-Myc protein. A., Q-MOL allosteric sites predictions using structures of individual amino acids as docking probes (see Methods). c-Myc nHLHZ domain (red) is shown in complex with Max (green), PDB 1NKP. **B., Q-MOL VLS results.** The docking site was defined around predicted allosteric site, adjacent to sequence stretch L943-KKATAYILS-V953 (indicated with a yellow circle, amino acid numbering corresponds to that of PDB 1NKP). The ranking of hits, listed in Table 3, is indicated with green circles. The complete human metabolites library was used for docking^46^. The top 235 out of 9000 hits are shown.

**Table 3.**
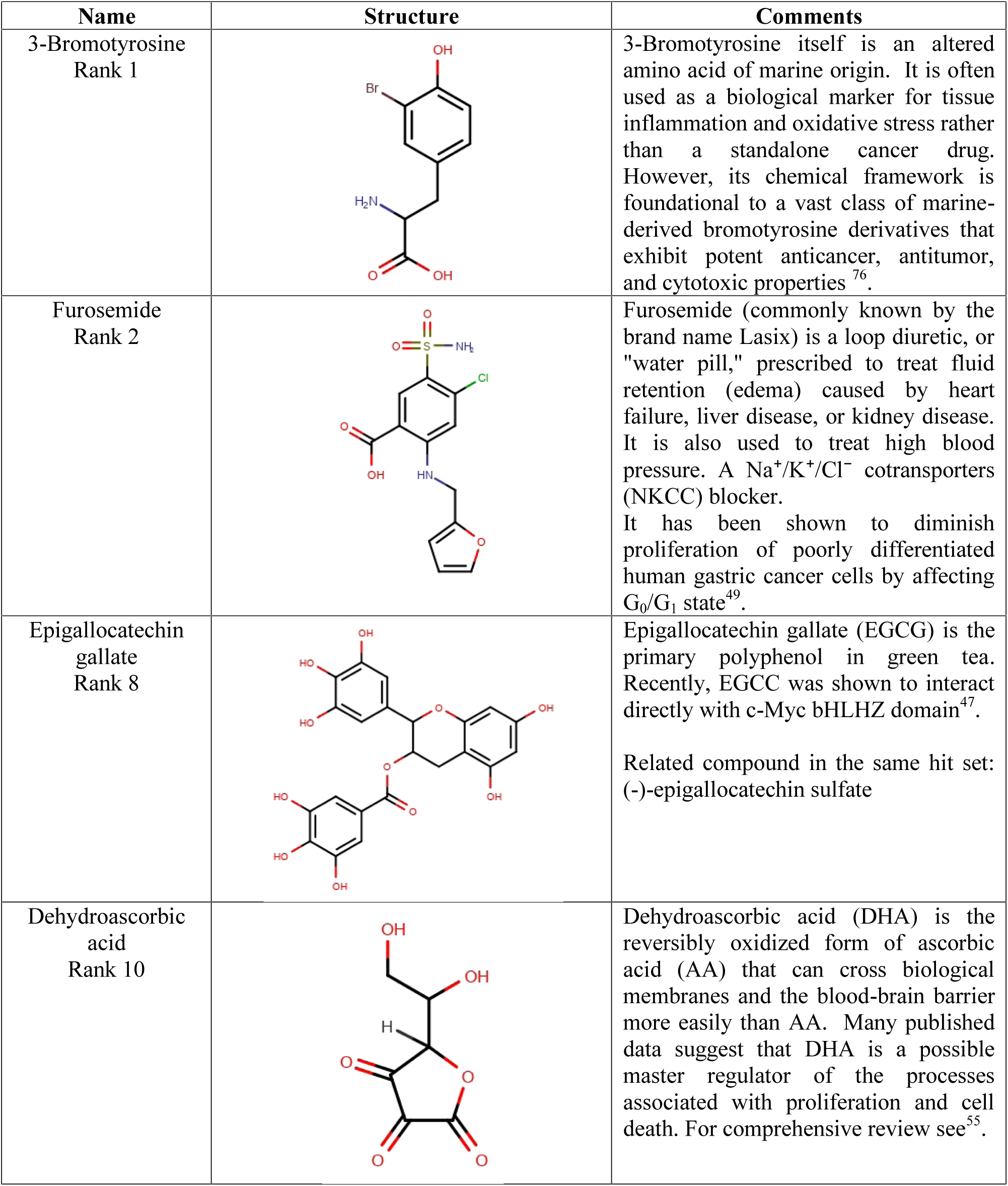
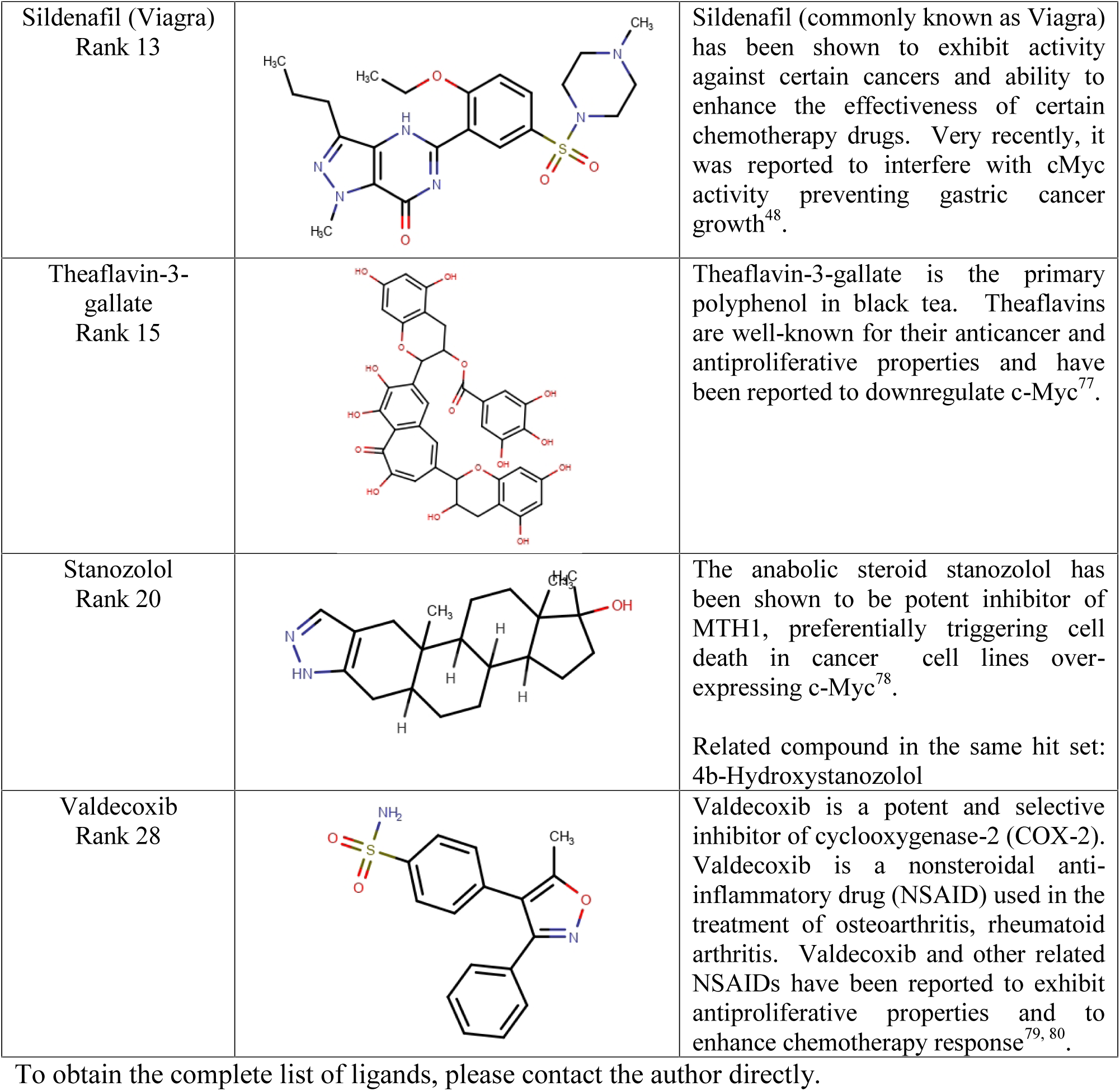
Selected annotated ligands from Human Metabolites library identified as hits in Q-MOL VLS against bHLHZ domain of c-Myc.

| Name | Structure | Comments |
| --- | --- | --- |
| 3-Bromotyrosine<br>Rank 1 |  | 3-Bromotyrosine itself is an altered amino acid of marine origin. It is often used as a biological marker for tissue inflammation and oxidative stress rather than a standalone cancer drug. However, its chemical framework is foundational to a vast class of marine-derived bromotyrosine derivatives that exhibit potent anticancer, antitumor, and cytotoxic properties<sup>76</sup>. |
| Furosemide<br>Rank 2 |  | Furosemide (commonly known by the brand name Lasix) is a loop diuretic, or "water pill," prescribed to treat fluid retention (edema) caused by heart failure, liver disease, or kidney disease. It is also used to treat high blood pressure. A Na<sup>+</sup>/K<sup>+</sup>/Cl<sup>-</sup> cotransporters (NKCC) blocker. It has been shown to diminish proliferation of poorly differentiated human gastric cancer cells by affecting G<sub>0</sub>/G<sub>1</sub> state<sup>49</sup>. |
| Epigallocatechin gallate<br>Rank 8 |  | Epigallocatechin gallate (EGCG) is the primary polyphenol in green tea. Recently, EGCC was shown to interact directly with c-Myc bHLHZ domain<sup>47</sup>.<br>Related compound in the same hit set: (-)-epigallocatechin sulfate |
| Dehydroascorbic acid<br>Rank 10 |  | Dehydroascorbic acid (DHA) is the reversibly oxidized form of ascorbic acid (AA) that can cross biological membranes and the blood-brain barrier more easily than AA. Many published data suggest that DHA is a possible master regulator of the processes associated with proliferation and cell death. For comprehensive review see<sup>55</sup>. |
| Sildenafil (Viagra)<br>Rank 13 |  | Sildenafil (commonly known as Viagra) has been shown to exhibit activity against certain cancers and ability to enhance the effectiveness of certain chemotherapy drugs. Very recently, it was reported to interfere with cMyc activity preventing gastric cancer growth<sup>48</sup>. |
| Theaflavin-3-gallate<br>Rank 15 |  | Theaflavin-3-gallate is the primary polyphenol in black tea. Theaflavins are well-known for their anticancer and antiproliferative properties and have been reported to downregulate c-Myc<sup>77</sup>. |
| Stanozolol<br>Rank 20 |  | The anabolic steroid stanozolol has been shown to be potent inhibitor of MTH1, preferentially triggering cell death in cancer cell lines over-expressing c-Myc<sup>78</sup>.<br>Related compound in the same hit set:<br>4b-Hydroxystanozolol |
| Valdecoxib<br>Rank 28 |  | Valdecoxib is a potent and selective inhibitor of cyclooxygenase-2 (COX-2). Valdecoxib is a nonsteroidal anti-inflammatory drug (NSAID) used in the treatment of osteoarthritis, rheumatoid arthritis. Valdecoxib and other related NSAIDs have been reported to exhibit antiproliferative properties and to enhance chemotherapy response<sup>79, 80</sup>. |

To obtain the complete list of ligands, please contact the author directly.

As a result of protein-ligand docking simulation, a set of 8 compounds (out of 9,000 screened ligands) were selected from top-ranking hits and annotated in Table 3, which highlights the most interesting candidates. For instance, epigallocatechin gallate (EGCG) (rank 8) is the primary antioxidant polyphenol in green tea that possesses potent anticancer properties. Recently, it has been shown to directly bind bHLHZ domain of c-Myc^47^. A related compound, (-)-epigallocatechin sulfate was also predicted as a the c-Myc binder with a similar rank.

Two FDA approved drugs were predicted as direct c-Myc binders: sildenafil (Viagra) (rank 13) and furosemide (Lasix) (rank 2). Sildenafil has been reported to interfere with c-Myc activity preventing human gastric cancer growth^48^, while furosemide has been shown to inhibit proliferation of human gastric cancer cells^49^. Note the gastric cancer cell type that correlates with each compound’s activity.

Finally, the most notable predicted hit - dehydroascorbic acid (DHA) (ranking 11 out of 9,000 metabolites). DHA is the major reversibly oxidized form of ascorbic acid (AA). DHA is actively imported into the endoplasmic reticulum (ER) of cells via glucose transporters^50^. Within ER, it is then reduced back to AA by glutathione and other thiols^51^. Importantly, the glucose transporters (e.g. GLUT1) transport DHA in most cells^52^, while transport of AA is limited to sodium-dependent transporters in specialized cells. AA is well known for its anticancer properties at high concentrations. The use of AA in cancer therapy remains a topic of considerable debate, numerous studies have investigated its potential anti-tumor effects, which are mostly attributed to its pro-oxidative activity within cancer cells and the resulting selective pressure in malignant tissue^53, 54^. DHA, on the other hand, is somewhat enigmatic compound, and only recently, its cell cycle regulatory and antiproliferative properties have been noticed^55^. Importantly, DHA has been reported to show significant cytotoxicity toward cancer stem cells (CSCs)^54^, while c-Myc has been shown to play a significant role in CSCs signaling and survival and to confer resistance to cancer therapy^56^. CSCs predominantly maintain a quiescent metabolic state but can dynamically shift between glycolytic and oxidative phosphorylation–dependent phenotypes, making their metabolism a challenging and unreliable therapeutic target^57^. In contrast, the *in silico* prediction that DHA modulates c-Myc activity by directly interacting with its transactivation domain positions DHA as a novel therapeutic agent capable of specifically targeting CSC-derived, treatment-refractory cancers. This direct binding also calls into question the currently accepted pro-oxidative mechanism proposed for the anticancer action of AA, suggesting instead a redox-independent, target-specific mode of action.

## Conclusion

The Q-MOL platform validation results presented here are just a small subset of protein targets processed using the Q-MOL software. The Q-MOL platform is the first of its kind to computationally implement principles of the energy landscape protein folding theory as applied to the problem of protein-ligand docking. To this end, Q-MOL implements a ligand-centric protein-ligand docking algorithm, that implicitly evaluates receptor flexibility using receptor representation as a parametric hyperspace of binding energy values, computed using re-parameterized OPLS force field. As expected from the energy landscape theory of protein folding, the progress through different computational phases of *in silico* drug discovery is tightly coupled to biological validation of intermediate hits. It has also been shown that Q-MOL ligand docking force field (based on OPLS force field) as well as ligand docking methodologies can be applied "As Is" to non-protein receptor targets such as non-coding viral RNA molecules. The independence of the docking simulation from the type of a target demonstrates the universal nature of Q-MOL ELT implementation and parameterization of ligand docking force field.

The Q-MOL software successes is proof that, contrary to a conventional belief, highly flexible proteins are significantly more amenable targets for drug discovery than rigid proteins such as enzymes. Flexible and intrinsically disordered proteins exist in a vast array of conformational states allowing for a significantly increased set of diverse ligand chemotype hits relative to that of rigid protein targets. The correct evaluation of the contribution of many possible conformations of a flexible protein target during protein-ligand docking simulation allows for discovery of specific and potent small molecule ligands that also modulate its function depending on a specific environment.

## Methods

### Q-MOL Primary Virtual Ligand Screening

The term *primary* refers to a VLS performed with a diverse unfocused chemical library, and the resulting hit set containing ligands predicted against all possible protein conformations. In case of a flexible protein target, whose apparent native state is environment-dependent (e.g. on a cell type), the primary hits set is re-validated using corresponding functional assays (Fig. 12).

The preparation of protein molecule includes adding of hydrogen atoms and the assignment of the standard Optimized Potential for Liquid Simulations (OPLS) ^58^ atom types. The location of a docking site and simulation boundary conditions are defined using convenient 3D graphics interface (Fig. 10A). The ligands are docked using the Markov Chain Monte Carlo simulation in the internal coordinate space as implemented in the Q-MOL program. To increase the speed of calculations, a subset of protein-ligand docking parameters is treated as grid-based potentials accounting for protein–ligand interactions. The preparation of each ligand for docking simulation initially includes an automatic OPLS atom type assignment and conversion of the 2D sketch-like models (input as the MDL MOL format of SDF file) into the 3D molecular models. The full-atom ligand structure is then minimized using the Q-MOL small molecule minimization protocol in OPLS force field. The small molecule structure optimization protocol combines minimization in both internal and Cartesian coordinates to properly optimize the rotable bonds of a small molecule. During the docking of a compound database, the compounds are minimally filtered by applying lower molecular mass cut-off of 200 Da, a polyphenols sub-structure filter, and a filter detecting chlorine atoms attached to aliphatic carbons. To increase the overall performance of VLS, the compound database is converted into a binary space partitioning (BSP) tree-like structure based on the chemical similarity between compounds (see *Compound Library Preparation*). Within the chemical BSP tree, the ligands are clustered by the chemical fingerprint similarity as implemented in the Q-MOL program. The chemical BSP tree is used by the Q-MOL VLS protocol for the efficient ligand database traversal during the VLS experiments. Because of the stochastic nature of the Q-MOL docking protocol, each ligand is docked at least three times, and the best (lowest) energy conformers is selected. To differentiate between the true and false binders, Q-MOL VLS uses a proprietary protein-ligand binding energy evaluation function. This function is based on the re-parameterized OPLS force field, and, in addition to protein-ligand interactions, accounts for the internal energy change of the docked ligand. Q-MOL protein-ligand methodology does not employ a trained ligand scoring function for hit ranking. The full protein flexibility treatment is implicitly addressed in the protein-ligand docking simulation by visiting all possible protein conformations in the parametric hyperspace in proximity to a selected docking site. Implicit treatment of protein flexibility during protein-ligand docking simulation is based on a computational variant of energy landscape theory (ELT) of protein folding (see *Results and Discussion* for details). Because the Q-MOL VLS docking curve represents predicted hits against many possible protein conformation, all of the predicted hits are treated as *equiprobable* protein target binders and are selected for biological evaluation across the whole docking curve. The exact implementation of the protocol will be described elsewhere. *Compound Library Preparation* To increase the overall performance of VLS, a compound library in converted into a binary space partitioning (BSP) tree-like structure (Fig. 10B). Within the chemical BSP tree, the ligands are clustered pairwise (hence binary) by the chemical fingerprint similarity as implemented in the Q-MOL program. The chemical BSP tree is used by the Q-MOL VLS protocol for the efficient ligand library traversal during the VLS experiments. During the traversal only focused ligand subsets (tree branches) of current best (most probable) binders are docked, while the chemical subsets of least probable binders are omitted. This technique allows screening of ligand libraries, containing millions of compounds, on a time scale of several days using ordinary desktop computers or cloud-based virtual machines instead of dedicated supercomputers. This feature becomes especially useful when docking virtual combinatorial libraries. These libraries can contain hundreds of millions of compounds.

### Computational Optimization of Primary VLS Hits

The primary VLS hits are biologically validated ligands, predicted after the very first round of VLS when a complete compound library was used for the docking. Because each of the primary hits targets diverse protein conformations in vicinity of some local minima, further computational optimization is necessary to converge on ligands targeting near apparent native receptor conformations. To this end, the chemical space of each primary hit is "extended" by searching for analogues in SDF compound library. The similarity between structures is measured by computing distance between proprietary Q-MOL chemical fingerprints which are based on protein-ligand docking parameters. For each hit, closest 256 analogues are retrieved, and combined together into a single ligands pool. All of the new ligands are than docked and sorted by relative binding energy. Because these ligands represent a structural subset that targets a conformational ensemble near apparent native state of a receptor, the top (lowest energy) hits are than selected for biological evaluation.

### Detection of Allosteric Ligand Binding Sites

The correct definition of ligand docking site for VLS experiments is one of the most critical tasks of *in silico* drug discovery methodology. The definition of the ligand docking site includes the starting point of the protein-ligand docking simulation, and the boundary conditions that limit the execution of protein-ligand docking simulation to some, proximal to the starting point, regions of protein target. Flexible non-enzyme protein targets, IDPs or flexible enzyme regulatory domains contain hidden, largely unannotated, ligand binding sites. These allosteric ligand binding sites usually have no classical binding pocket features, and the actual binding pockets are formed during a specific structural transaction when a receptor binds a ligand, and potentially other additional binding partners. Thus, a computational tool for the protein target annotation is a prerequisite for the successful *in silico* drug discovery campaign. To this end, the Q-MOL drug discovery platform implements a unique methodology that allows for reliable prediction of such allosteric ligand binding sites on a surface of a receptor. The methodology is based on the thermodynamic fact that efficient binding occurs only when the net free energy change is negative. There are many sources contributing to the energy minimization during a binding event, one the primary sources is associated with protein molecule itself, and specifically with the actual ligand binding site. This excess free energy is stored within a protein surface mainly as a free energy of unrealized non-bonded interactions and non-minimized torsional strains. To detect this energy excess, the QMOL ligand binding sites detection method applies exactly the same protein-ligand docking algorithm as the one used for VLS. Briefly, the molecular surface of a receptor is systematically "scanned" by docking "probe" ligands at evenly distributed locations across the surface. During the docking of a probe ligand, the best (lowest) predicted binding energy values is collected together with coordinates of the corresponding probe conformer. Because of the stochastic nature of a docking simulation, the systematic surface scan is ran at least 3 times. The collected binding energy values are then normalized and converted into probabilities of binding of probe ligands at specific binding sites (Fig. 7, 8, 11A). Finally, a *specificity index* parameter is computed for a probe ligand (Fig. 11B). Specificity index reflects preference of binding of a probe ligand toward a specific spot on the molecular surface of a protein. When a probe ligand displays a preference to a particular binding site, high probability values of binding tend to cluster in space in proximity to that binding site. This clustering trend is quantified as a specificity index. The specificity index is a relative value, and it is computed within a probe set. The Q-MOL ligand binding sites detection tool has two main application: receptor surface scanning by known small molecule binders (Fig. 7), and receptor surface scanning by 20 individual amino acids represented as small molecule ligands (Fig. 8, 11). The first application is used to locate a putative ligand binding site and/or validate a known ligand, the latter is used for annotation of protein targets in the very beginning of a drug discovery campaign.

### Detection of Allosteric Binding Sites on the Surface of Non-coding Viral RNA Molecules and RNA-ligand Docking

All of the applicable Q-MOL methodologies were used for RNA molecules "As Is" without any specific docking parameter or computational protocol modifications. To detect putative small molecule binding sites on the surface of RNA molecules, the molecular surface scanning protocol was applied using structures of 20 individual amino acids as docking probes. The most probable small molecules binding sites were then selected for RNA-ligand docking simulation setup. Essentially, the RNA molecules were treated exactly the same way as protein molecules in terms of ligand docking simulation setup, computational protocols and their parameters.

### Additional Q-MOL Software Features and Architecture

The OPLS all-atom force field is uniformly used through software ensuring coherent energy calculations across the boundaries of distinct computational tools (protein structure optimization, protein-ligand docking, small molecule minimization, chemical fingerprints) ^58^. For efficient parameters sampling and minimization, Q-MOL renders structures of both proteins and small molecules in internal coordinates. The representation of molecular structures in internal coordinates was introduced by Scheraga H.A. in 1968 ^59^. The versatility of internal coordinates was further asserted by ^60^ and ^61^. Markov Chain Monte Carlo (MCMC) algorithm is used throughout the software to efficiently converge to energy minimum in the course of multiparametric simulations in internal coordinates. The essential cheminformatics tools based on SMILES/SMARTS presentation of small molecules are implemented. These tools are used in automatic OPLS atom type assignment, compounds search by substructure or Q-MOL chemical fingerprint. A user enjoys intuitive goal-oriented UI. All heavy computing is seamlessly off-loaded to cloud compute servers so that there is no need to provide specialized on-site computing infrastructure.

## Acknowledgment

The author would like to express sincere gratitude to Dr. Wayne Guida (University of South Florida) for taking the time and effort necessary to review the manuscript.

## Data and Software Availability

The Q-MOL software code was written solely by the author. The code is proprietary and cannot be disclosed at this point. The certain number of Q-MOL licenses can be made available for academic organizations and interested readers free of charge. Additionally, several Q-MOL features are available free of charge via the website https://q-mol.org. The author believes that publishing this manuscript is important for the field of *in silico* drug discovery because it successfully addresses several rather controversial molecular modeling problems with concrete prior experimental validation. Please enquire with Dr. Anton Cheltsov for information about obtaining a license, or the website account credentials.

